# Faf1 accelerates p97-mediated protein unfolding by promoting ubiquitin engagement

**DOI:** 10.1101/2025.10.27.684972

**Authors:** Zengwei Liao, Connor Arkinson, Andreas Martin

## Abstract

P97/VCP is a protein unfoldase of the AAA+ ATPase family that plays essential roles in numerous processes, including ER-associated degradation and DNA replication. For unfolding of proteins that are modified with K48-linked ubiquitin chains, p97 works with the heterodimeric cofactor Ufd1-Npl4, and the cofactor Faf1 was shown to enhance this activity during replisome disassembly, yet the mechanisms remain unknown.

Here, we employ an in vitro reconstituted system with human components for biochemical experiments, FRET-based assays, and cryo-EM structure determination to reveal that Faf1 plays a generic role in accelerating ubiquitin-dependent substrate processing by promoting the unfolding of an initiator ubiquitin and its engagement by the ATPase. Faf1 thereby uses its p97-bound C-terminal UBX domain to anchor a long helix that braces the UT3 domain of Ufd1 and stabilizes the Ufd1-Npl4 cofactor for ubiquitin unfolding. Our findings demonstrate how p97 works simultaneously with several cofactors to facilitate the unfolding of ubiquitinated proteins, indicating more complex regulatory mechanisms for substrate selection than for the simpler yeast Cdc48.

## Introduction

The mammalian Valosin-Containing Protein (VCP) or p97 (Cdc48 in yeast) is an essential AAA+ (ATPases Associated with various cellular Activities) protein unfoldase that plays roles in many cellular processes, such as endoplasmic reticulum-associated degradation (ERAD), endosomal trafficking, membrane fusion, DNA damage response, and DNA replication ^1–6^. A major function of p97 is to extract proteins from macromolecular complexes or membranes and unfold them prior to degradation by the 26S proteasome^7^. Many of p97’s substrates are therefore marked with polyubiquitin chains. Mutations in p97 are for instance linked to cardiovascular and neurodegenerative diseases, including multisystem proteinopathy (MSP), a fatal disease with various clinical pathologies that affect the brain, muscle, and bone^8–10^.

P97 is a homo-hexamer with each protomer containing a N-terminal domain (NTD) and tandem ATPase domains, D1 and D2, that in the hexamer form stacked rings. Many cofactor proteins bind p97 to contribute various activities or regulate its function, including substrate recruitment, engagement, unfolding, and deubiquitination^11^. Cofactors thereby use ubiquitin regulatory X (UBX/UBXL) domains or linear sequences such as VCP-binding motifs (VBMs), VCP-interacting motifs (VIMs), or SHP motifs (also called BS1, binding segment 1) to interact with p97’s NTD, while others bind the C-terminal tail of p97 through a PUB (peptide:N-glycanase and UBA or UBX-containing proteins) or PUL (PLAP, Ufd3p, and Lub1p) domains. One of the most well studied p97 cofactors is the heterodimeric Ufd1-Npl4 (UN) complex, which binds on top of the p97 hexamer, recruits ubiquitinated substrates, and initiates their unfolding, with specificity for K48-linked ubiquitin chains^12,13^. Npl4 interacts through its UBXL and zinc-finger domains with p97’s NTD and D1 domains, while Ufd1 contacts Npl4 through a small interface and is highly flexible, which so far prevented its visualization by cryo-EM^12^. Structural and functional analyses of UN-bound Cdc48 indicate that an initiator ubiquitin in the substrate-attached ubiquitin chain gets unfolded by UN in an ATP independent manner and binds in a hydrophobic groove on Npl4 to insert its N-terminus into the Cdc48 central channel. The Cdc48 motor then utilizes ATP hydrolysis to engage and pull on the initiator ubiquitin, unfold and translocate any proximal ubiquitin moieties in the chain, and thread the attached substrate protein itself^12,14^. The ubiquitin moiety immediately distal to the initiator was observed to interact with parts of Npl4 and Ufd1, while the consecutive next two ubiquitins bind on top of the so-called tower domain of Npl4^12,15^ and increase substrate affinity during recruitment^14^. Importantly, the ubiquitin moiety on the proximal, substrate-facing side of the initiator binds Ufd1’s UT3 domain, which appears to determine the K48-ubiquitin linkage specificity of Cdc48-UN and is critical for insertion of the initiator ubiquitin into the ATPase motor^14^.

Interestingly, only the proximal ubiquitin(s) and the substrate appear to get unfolded and threaded, whereas ubiquitins on the distal side of the initiator may not be unfolded^14,16–18^. Release of the ubiquitinated substrate from the motor after unfolding therefore represents the rate-limiting step for both Cdc48-UN and p97-UN, at least in the absence of other cofactors or deubiquitinases^14,18^. Furthermore, without any additional cofactors, p97-UN is far less efficient in substrate unfolding than Cdc48-UN and requires much longer ubiquitin chains^19,20^, suggesting more complex regulatory mechanisms for substrate recruitment and processing. In humans there is an expanded family of p97 cofactors, but it remains unclear how, when, or which of these cofactors work together. Many of them may act independently, compete for the same binding sites on p97, or function only in the context of specific substrates. However, recent insights into replisome disassembly showed that fas-associated factor 1 (Faf1), works with p97-UN to enhance the unfolding of MCM complexes and their removal from DNA, potentially through lowering the threshold for the ubiquitin-chain length^19^. Interestingly, in addition to replisome disassembly^19,21^, Faf1 has been reported to play significant roles in other pathways, including immune response^22,23^ and aggresome clearance^24^, indicating possible generic roles in multiple p97-dependent processes.

Faf1 is a multidomain protein (Fig. 1A) that uses its C-terminal UBX domain for binding to p97’s NTD^25^, potentially interacts with ubiquitin chains through its N-terminal UBA (ubiquitin-associated) domain^26^, and contacts other chaperones, like Hsp70, through one of its two UBL domains^27^. However, the mechanism for Faf1-mediated enhancement of substrate unfolding by p97-UN remains unclear.

**Figure 1.**
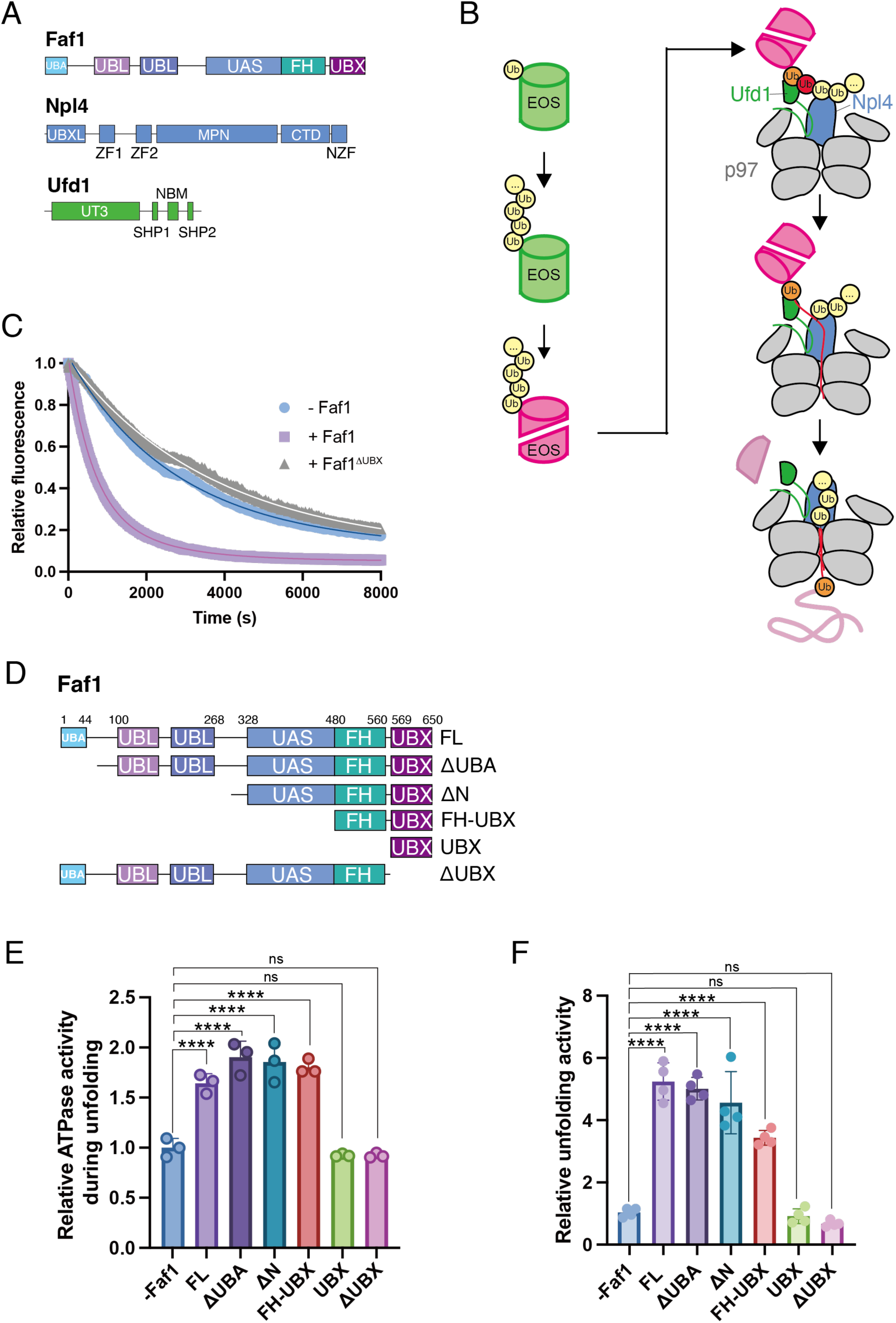
Faf1 accelerates the unfolding of ubiquitinated substrates. **A)** Schematics of domain architectures for human Faf1, Npl4, and Ufd1. **B)** Left: Overview of substrate preparation. mEos3.2 fused to mono-ubiquitin was poly-ubiquitinated and photoconverted by UV irradiation to red Eos with cleaved backbone. Right: The substrate-unfolding assay monitors the loss of red Eos fluorescence. An initiator ubiquitin (red sphere), the proximally neighboring ubiquitin (orange sphere), and ubiquitins on the distal side (yellow spheres) bind to the UT3 domain of Ufd1 (green) and the top of Npl4 (blue), before the initiator ubiquitin is unfolded, captured by Npl4, and inserted into p97, followed by unfolding and threading of the entire proximal part of the chain as well as the Eos substrate. Ubiquitin moieties located on the distal side of the initiator do not get unfolded by p97. **C)** Traces for the loss of red Eos fluorescence upon unfolding by p97-UN in the absence and presence of Faf1. All traces represent the averages of three independent measurements that were normalized to the starting fluorescence value. Solid lines show the single-exponential fits. **D)** Schematics of Faf1 truncation mutants used in biochemical experiments. **E)** Relative ATPase activities of p97-UN during unfolding of ubiquitinated green Eos in the absence of Faf1 (normalized to 1) and the presence of full-length or truncated Faf1. Shown are the mean values and standard deviations of the mean for three technical replicates. Statistical significance was calculated using a one-way ANOVA test: ****p<0.0001; ns, p>0.05. **F)** Relative substrate-unfolding activities of p97-UN in the absence of Faf1 (normalized to 1) and in the presence of saturating (2 μM) full-length or truncated Faf1. Shown are the mean values and standard deviations of the mean for four technical replicates. Statistical significance was calculated using a one-way ANOVA test: ****p<0.0001; ns, p>0.05.

Here, we *in vitro* reconstitute p97-UN unfolding of model substrates with K48-linked ubiquitin chains and reveal that Faf1 has a generic function by facilitating initiation, increasing substrate capture, and thus accelerating processing. Our FRET-based studies show that Faf1 specifically accelerates initiator-ubiquitin insertion into p97, with rates that are comparable to those of Cdc48-UN. Interestingly, p97 subsequently unravels the mEos model substrate much more slowly than Cdc48, suggesting mechanical unfolding as the rate-limiting step for p97. Using cryo-EM single-particle analysis, we observe the p97-UN-Faf1 complex with a ubiquitin chain during unfolding initiation. Faf1’s C-terminal UBX domain binds to the NTD of p97 and stabilizes the SHP motif of Ufd1, which is key to Faf1 recruitment. A helix preceding the UBX domain reaches up and braces Ufd1’s UT3 domain. This UT3 domain shows at least two bound ubiquitin moieties, with one being linked to the unfolded initiator ubiquitin that interacts with Npl4’s hydrophobic groove and reaches into the p97 hexamer. Site-specific mutagenesis combined with our substrate-unfolding and FRET-based assays confirmed that Faf1’s helix interactions with the UT3 domain of Ufd1 allow for more efficient initiator-ubiquitin unfolding and insertion into p97. Finally, we show that this mechanism is conserved for the membrane associated cofactor Faf2, which plays critical roles in ERAD^28,29^. Faf1 and Faf2 promote a more productive complex between UN, p97, and ubiquitinated substrates that allows higher efficiency in ubiquitin insertion and unfolding initiation, which indicates how those cofactors may generically support the numerous cellular functions of p97-UN.

## Results

### Faf1 stimulates unfolding activity of p97-UN

To test whether Faf1 can generally accelerate the unfolding of ubiquitinated substrates by p97-UN, we utilized a photoconverted, red-fluorescent and thus irreversibly unfolding mEos3.2 model substrate with a covalently fused mono-ubiquitin that was enzymatically extended into long (> 8 moieties) K48-linked polyubiquitin chains (Fig. 1B, Supp. Fig. 1A,B)^30,31^. Human p97, UN, and Faf1 were recombinantly expressed and purified from *E.coli*. Reconstituted Eos-unfolding experiments under single-turnover conditions (i.e. enzyme at saturating concentrations and in excess over substrate) in the absence or presence of Faf1 showed that Faf1 accelerates p97-UN-mediated substrate processing by ∼ 5-fold, from 1_Eos unfold_ (-Faf1) = 4727 ± 688 s to 1_Eos unfold_ (+Faf1) = 902 ± 76 s (Fig. 1C; Supp. Fig. 1C). These results are overall consistent with previous findings about the Faf1 effects on MCM-complex disassembly^19^, but, importantly, also demonstrate a role in accelerating unfolding of ubiquitinated substrates in general.

### A helix-UBX domain fragment of Faf1 is sufficient for facilitating substrate unfolding

To assess which parts of the multi-domain Faf1 are responsible for the stimulation of substrate unfolding by p97, we generated several truncated variants (Fig. 1D, Supp. Fig. 1D) and tested their effect on both substrate processing and ATP hydrolysis (Fig. 1E,F). Faf1 alone, in the absence of UN and substrate, did not increase p97’s low basal ATPase activity (Supp. Fig. 1E). However, Faf1 stimulated ATP hydrolysis ∼ 1.7-fold in the presence of substrate and UN (Fig. 1E), suggesting that it either accelerates the motor during threading and unfolding or it facilitates substrate engagement, thereby increasing the fraction of actively substrate-processing p97 with ATP hydrolysis above idling^18^. While Faf1’s UBA and UBL domains were dispensable for p97 activation, deletion of its UBX domain abolished Faf1’s stimulatory effects (Fig. 1E,F), which is consistent with previous observations for p97-mediated replisome disassembly in the presence of Faf1^19^. Faf1’s two UBL domains were suggested to function in binding other chaperones, such as Hsp70, and thus may play context-specific roles^23,27^, similar to the UAS domain that appears to bind fatty acids ^32^. We found that a minimal C-terminal fragment consisting of the FH and UBX domain (FH-UBX) leads to a stimulation of p97’s unfolding activity that is comparable to that of full-length Faf1, whereas the UBX domain alone was insufficient for stimulation (Fig. 1F).

### Faf1 does not enhance the activity of p97 bound to yeast UN

Since there are clear differences between human and yeast UN, we wondered whether human UN is more dynamic and therefore depends on Faf1 to form a productive complex for unfolding initiation. Interestingly, yeast UN (yUN) can support p97 unfolding of the ubiquitinated-Eos substrate at a rate of 1_Eos_ = 26.7 ± 2.4 min, which is ∼ 3.5-fold higher than the rate in the presence of human UN (hUN) without Faf1 (Supp. Fig. 1F) and demonstrates the functional conservation of the AAA+ motors. The addition of full-length Faf1 did not further stimulate the unfolding activity of p97-yUN, but in fact reduced it (Supp. Fig. 1F), potentially because Faf1’s UBL domains bind to the tower ubiquitin-binding sites of the yeast Npl4. We therefore used the FH-UBX domain fragment of Faf1 and observed an unfolding activity that was comparable to that of p97-yUN (Supp. Fig. 1F). yUN may intrinsically encode for more productive ubiquitin unfolding and insertion into Cdc48 or p97, whereas hUN is potentially more dynamic and depends on additional interactions such as Faf1’s FH-UBX-domain portion.

### Cryo-EM reveals a pre-initiation state with flexibly bound Npl4

For gaining structural insights into Faf1’s stimulating effects on p97 unfolding activity, we aimed to capture a substrate-engaged p97-UN-Faf1 complex. In a first attempt, we initiated the unfolding reaction with ATP, then switched through buffer-exchange to the non-hydrolyzable analog ATPγS in order to stall substrate-engaged complexes (Supp. Fig. 2A,B), and analyzed the sample by cryo-EM (Supp. Fig. 3). However, single-particle analysis resolved only the p97 hexamers with no defined density for most of Faf1, the UN complex, or the substrate, likely due to high flexibility and conformational heterogeneity. Symmetry expansion and focused classification of individual p97 promoters with a mask on the NTD resolved Faf1’s UBX domain bound to the NTD (Supp. Fig. 4A,B), which is in agreement with previous crystal structures^33,34^, and just parts of the preceding long helix extending up from the NTD.

We therefore turned to a different experimental setup that used p97-UN with the FH-UBX fragment of Faf1 and long unanchored ubiquitin chains instead of the ubiquitinated model substrate, in an attempt to reduce conformational heterogeneity. We reconstituted these components in the presence of ATP to promote functional complex states during unfolding initiation. Cryo-EM analysis of this sample revealed a minor fraction of particles in a potential apo or pre-initiation (PI) state, in which Npl4 is resolved on top of the p97 hexamer, but there is no indication of ubiquitin, and only the NTD-bound SHP and UBX domains are visible for Ufd1 and Faf1, respectively (Supp. Fig. 5-7). Npl4 is at lower resolution and apparently flexibly bound, using its UBX domain to interact with the NTD of protomer E and its zinc finger 1 (ZF1) to contact the D1 ATPase domain of protomer F in the p97 hexamer, while ZF2 does not interact with an ATPase domain and is unresolved (Supp. Fig. 7B). The planar D1 ring in this PI state has protomers A, B, and F occupied with ADP, and protomers C-E bound to ATP, whereas the entire D2 ring is bound to ADP (Supp. Fig. 7E), similar to what was observed for the D2 ring in Cdc48-UN during initiation^15^. In this state, the “Npl4 loop” (residues 425-439) is positioned right above the entrance to the p97 channel, in a conformation that would likely inhibit initiator-ubiquitin insertion into the motor (Supp. Fig. 7A). This also resembles the previously described ubiquitin-bound Cdc48-UN complex, in which the Npl4 loop adopted a similar position and ubiquitin was absent from the Npl4’s groove^15^. However, in contrast to our structure of human p97-UN, the yeast complex showed ubiquitins bound to the top of Npl4, which interacted through both ZF1 and ZF2 with the D1 ring of p97.

### Faf1 braces the UT3 domain of Ufd1 for initiator unfolding

The majority of particles in our dataset represented initiation-state (I state) complexes, with p97 bound to UN, Faf1, and a ubiquitin chain during motor insertion. Local refinement of the p97-Npl4 portion revealed Npl4 stably bound to the D1 ring, with ZF1 and ZF2 contacting protomers F and D, respectively (Supp. Fig. 7C). The D1 ring contains three ADP and three ATP, and thus has an identical nucleotide occupancy as in the pre-initiation state (Supp. Fig. 7E), but protomer E and F are noticeably shifted downward (Supp. Fig. 7D). The entire D2 ring is again filled with ADP (Supp. Fig. 7E). Similar to the initiation state previously observed for Cdc48-UN^12^, an unfolded initiator ubiquitin (Ub^ini^) is bound to Npl4’s groove and reaches with its N-terminus into the p97 central channel, with Lys11 as the last resolved residue positioned near the bottom of the D1 ATPase ring (Fig. 2A; Supp. Fig. 7D). Through further focused classification with a mask on Npl4, we were able to identify a sub-state that showed the three ubiquitin moieties linked to Ub^ini^ on its distal side, Ub^dist1^, Ub^dist2^, and Ub^dist3^, bound on top of Npl4 (Fig. 2B). While Ub^dist2^ is in a stable position, Ub^dist1^ shows higher heterogeneity between subclasses, and Ub^dist3^ is visualized in only a few classes, located in a different position compared to previous structures of the ubiquitin-bound yeast Cdc48-UN (Fig. 2B)^15^. We also observed two Faf1 UBX domains, bound to the NTDs of p97 protomers B and C, and additional density extending from these domains toward Npl4. To improve the resolution and provide insight into the ubiquitin chain on the proximal side of the Npl4-bound unfolded Ub^ini^, we used a mask around the NTD of protomer C, Faf1’s UBX-helix portion, and the space toward Npl4 for further 3D classifications (Fig. 3). Global refinement of these classes revealed improved density for the Faf1 helix on p97 protomer C and the interacting density, which was particularly prevalent in two classes that we name initiating conformation 1 and 2 (IC1 and IC2, Fig. 3A,B). For identifying this additional density in contact with the Faf1 helix, we used AlphaFold predictions of a p97-UN-Faf1-ubiquitin complex, which indicated direct interactions of the Faf1 helix with the UT3 domain of Ufd1 (Fig. 3A). Indeed, in IC1 we could model the structure of the UT3 domain with two bound ubiquitins into the extra density, and we name these ubiquitins Ub^prox^ and Ub*. The interaction sites of Ub^prox^ and Ub* on the UT3 domain were previously described in NMR-binding studies of polyubiquitin and monoubiquitin, respectively ^35^. Notably, the C-terminus of Ub^prox^ is not in proximity of Ub*’s K48, indicating that these ubiquitins are not linked, but it points toward an additional density that potentially represents a third, poorly resolved ubiquitin on the UT3 domain (Supp. Fig. 8).

**Figure 2:**
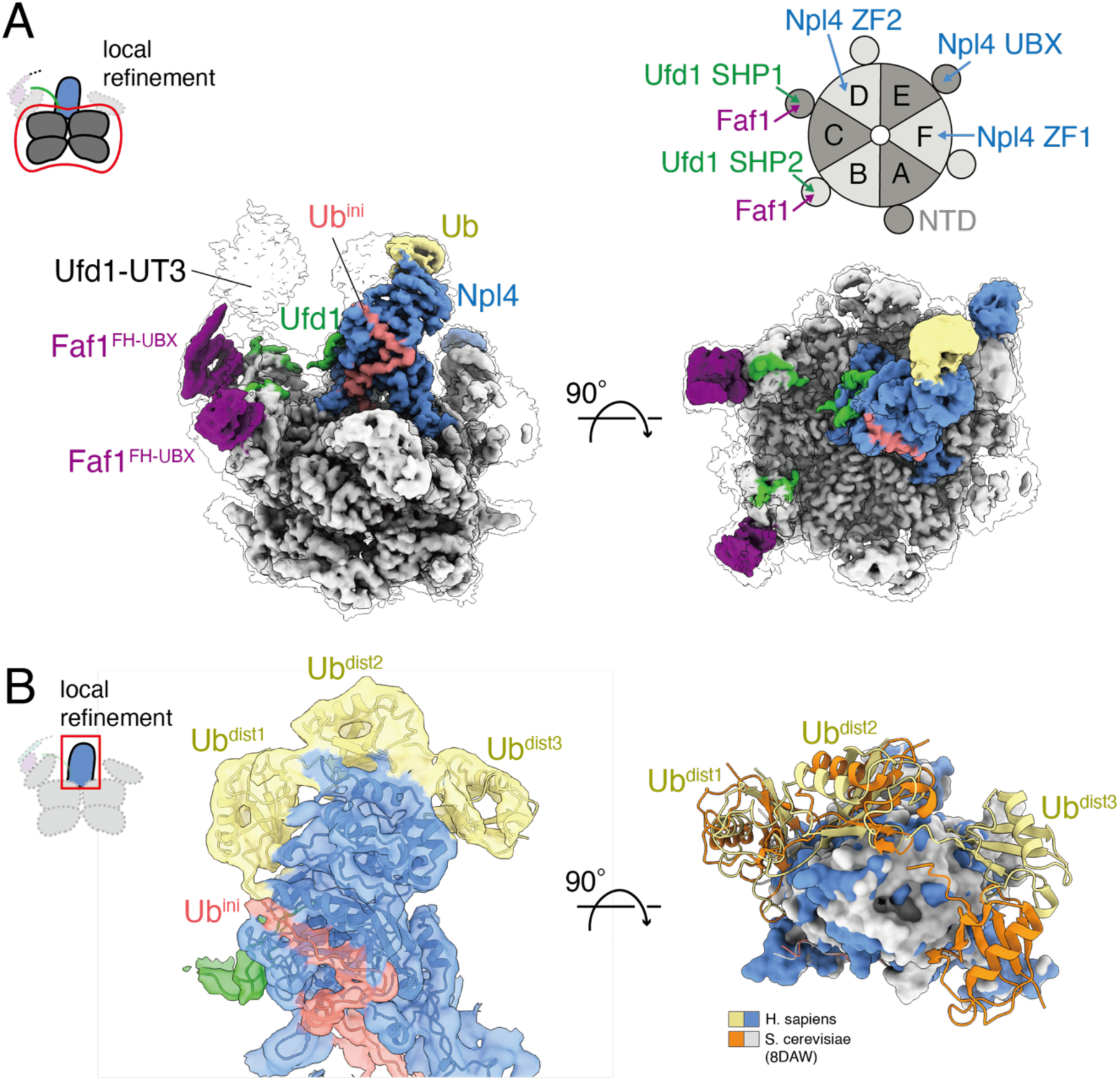
Cryo-EM structure of the poly-ubiquitin bound p97-UN-Faf1^FH-UBX^ initiation complex. **A)** Local refinement of the ATPase motor reveals Npl4 (blue) on top of the p97 hexamer (gray) with a bound initiator ubiquitin (Ub^ini^, salmon) reaching into the D1 ATPase ring. The SHP1 and SHP2 motifs of Ufd1 (green) occupy two neighboring NTDs of p97 together with two Faf1 UBX domains (purple). A lower threshold map is shown transparent to visualize Ufd1’s UT3 domain and extra density on Npl4. Top right: Schematic of the p97 hexamer with protomers A–F, illustrating the binding positions of Npl4 (ZF1; ZF2, UBX), Ufd1 (SHP1, SHP2), and Faf1 on the D1 ATPase ring. **B)** Left: Locally refined cryo-EM density focused on Npl4 and its interacting ubiquitins. Three distal ubiquitins (Ub^dist1^– Ub^dist3^, yellow) are resolved and bound to top of Npl4 (blue), with Ub^ini^ (salmon) in the hydrophobic groove and the Npl4-binding motif of Ufd1 (green) visible at the base. Right: Atomic model for the human Npl4 (blue surface) in complex with Ub^dist1^– Ub^dist3^ (yellow ribbons) overlayed with the structure of the ubiquitin-bound yeast Cdc48-UN (Npl4 as white surface, Ub^dist1^– Ub^dist3^ as orange ribbons; PDB ID: 8DAW).

**Figure 3:**
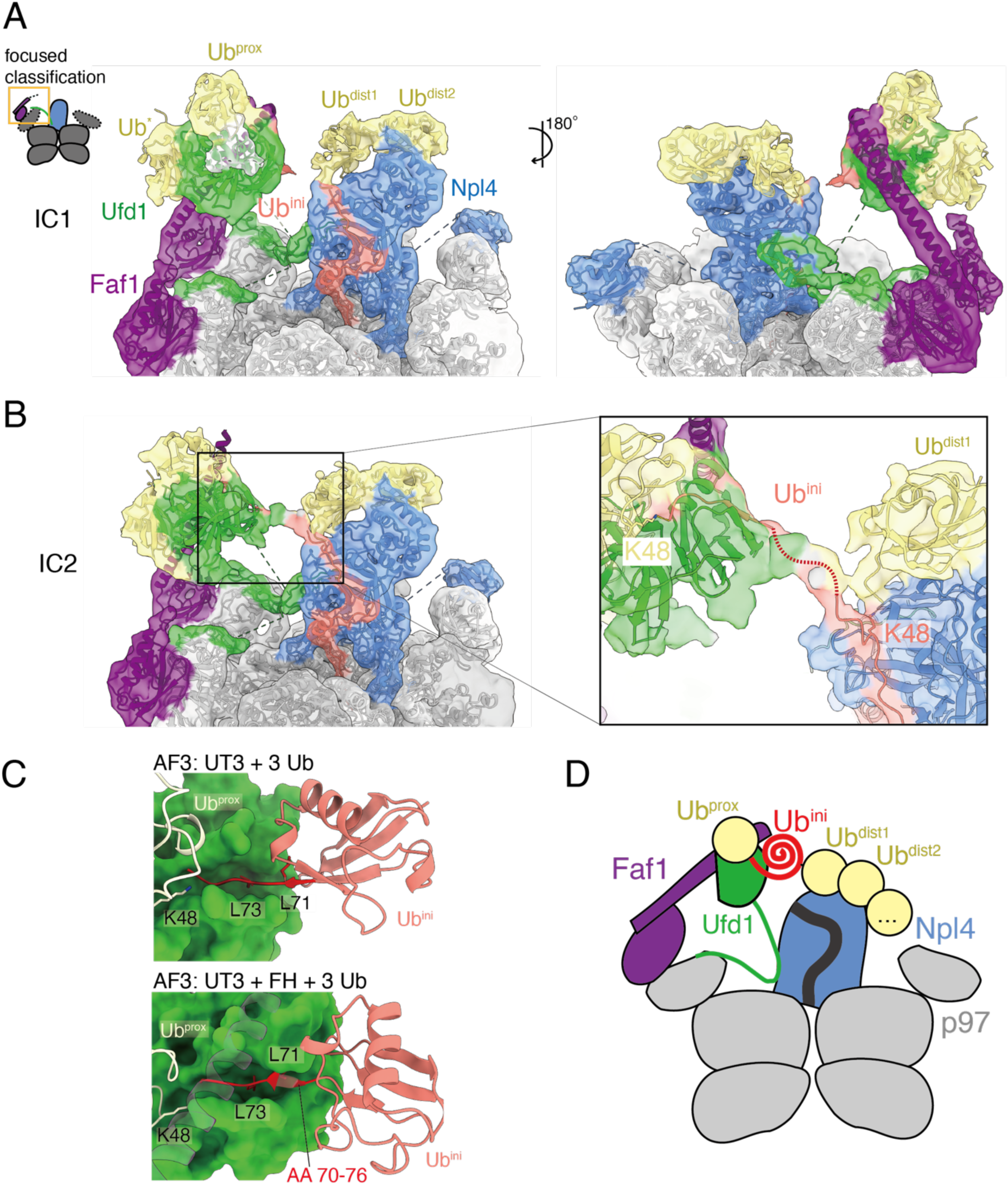
Faf1 braces the ubiquitin-bound UT3 domain of Ufd1 for initiation. **A)** Cryo-EM density map of the initiating conformation 1 (IC1) is shown in front and back view, with p97 in gray, Npl4 in blue, Ub^ini^ in salmon, the proximal and distal ubiquitins (Ub^prox^ and Ub^dist^) in yellow, Ufd1 in green, and Faf1^FH-UBX^ in purple. **B)** IC2 density map with the inset showing connecting density between the Npl4-bound Ub^ini^ and K48 of Ub^prox^, and a red dashed line indicating the potential path of the unfolded Ub^ini^ backbone. **C)** Top: AlphaFold model of Ufd1’s UT3 domain with three ubiquitins suggests an additional binding site accommodating the last seven residues (red) of Ub^ini^ (salmon). Bottom: Inclusion of Faf1’s FH in the AlphaFold predictions leads to a different conformation of Ub^ini^’s C-terminus, with residues 70-72 forming a small beta sheet with the UT3 domain and L71 as well as L73 docking into hydrophobic pockets. **D)** Schematic model for a putative pre-initiation state that precedes IC1 and IC2, in which Ub^dist^ moieties are bound to the top of Npl4, Ub^prox^ interacts with Ufd1’s UT3 domain, and Ub^ini^ is destabilized by binding of its C-terminal residues to UT3, mediated by Faf1.

In IC2, we also identified Ufd1’s UT3 domain with slightly less well-resolved Ub^prox^ and Ub* moieties, but noticed a shift in its position relative to Npl4 (Fig. 3B). Furthermore, there was continuous linking density from the UT3 domain to the unfolded Ub^ini^ in the Npl4 groove. Interestingly, AlphaFold models predict another ubiquitin binding site on the UT3 domain, in which ubiquitin’s G76 points toward K48 of Ub^prox^, the preceding C-terminal residues form a small beta-sheet with the UT3 domain, and Leu71, normally part of ubiquitin’s hydrophobic core, docks into a hydrophobic pocket of UT3 (Fig. 3C). We previously found for yeast Cdc48-UN that the ubiquitin moiety proximal to the initiator binds to the UT3 domain and that this interaction is responsible for the K48-linkage selectivity of ubiquitin unfolding by Cdc48-UN^14^. We therefore reasoned that the AlphaFold-predicted, C-terminally bound ubiquitin on the UT3 domain represents the initiator-ubiquitin moiety prior to its unfolding. Although the linking density between UT3 and Npl4 is too low in resolution to unambiguously model the C-terminus of Ub^ini^, it directly connects the Npl4-bound Ub^ini^ with K48 of Ub^prox^ on the UT3 domain, supporting our hypothesis that the C-terminus of the unfolded initiator ubiquitin binds to the UT3 domain in the AlphaFold-predicted position. This is also consistent with a recent study that investigated this binding site for ubiquitin’s C-terminal tail on the UT3 domain through X-ray crystallography and mutagenesis^36^.

In both IC1 and IC2, the FH helix of the Faf1 copy on the p97 protomer C contacts the UT3 domain and Ub^prox^ (Fig. 3). The direct contact between FH and ubiquitin at the UT3 domain is supported by photo-induced crosslinking experiments, in which we used Cy3-labeled ubiquitin chains and Cy5-labeled Faf1 FH-UBX fragment with benzophenylalanine (BPA) incorporated at position 501 to show that Faf1’s ubiquitin interaction depends on the presence of p97 and UN (Supp. Fig. 9). These findings are also consistent with recent other reports of interactions between FH and ubiquitin^37^. Faf1 likely serves as a stabilizing scaffold that bridges p97’s NTD with the UT3 domain, positioning UT3 to facilitate the unfolding of Ub^ini^, its capture by the Npl4 groove, and insertion into the p97 motor. Indeed, performing AlphaFold predictions of ubiquitin-UT3 interactions with and without FH suggests that the presence of the Faf1 helix promotes a different conformation of Ub^ini^’s last seven residues on the UT3 domain, which may have an effect on Ub^ini^ binding and unfolding (Fig. 3C). It is conceivable that in an initial binding stage the ubiquitin chain drapes over the p97-mounted UN cofactor, with Ub^prox^ and the C-terminus of the Ub^ini^ interacting with Ufd1’s UT3 domain, while the consecutive Ub^dist1^, Ub^dist2^, and Ub^dist3^ bind on top of Npl4 (Fig. 3D). Interaction of Ub^ini^’s C-terminus with the UT3 domain, facilitated by Faf1, is expected to destabilize Ub^ini^ and induce its unfolding for binding to the Npl4 groove.

Interestingly, from the UBX domain of the protomer B-bound second Faf1 we observed low-resolution density extending up to the opposite side of the UT3 domain, such that UT3 appears to be braced by the helices of two Faf1, possibly for additional stability or positional control (Supp Fig. 8A). Individual subclasses show the helix of the second Faf1 differentially resolved, depending on the movement of the UT3 domain relative to Npl4 (Supp. Fig. 8B), which further supports our model for Faf1-mediated UT3-domain positioning and a potential role in initiator-ubiquitin unfolding.

The number and location of Faf1 molecules on the p97 hexamer seems thereby determined by the SHP motifs of Ufd1. Using a locally refined map from our first cryo-EM data set together with AlphaFold predictions, we identified Ufd1’s SHP1 motif bound to the NTD of p97 protomer C, with its L235 anchored in a hydrophobic hole of the NTD (Fig. 3C,D; Supp. Fig. 10)^38,39^ and a linking density to Npl4, while the C-terminal SHP2 motif is bound to the NTD of protomer B (Fig. 3C,D). Although it cannot be ruled out that the presence of two Faf1 copies is a consequence of our *in vitro* reconstitution, the correlation between Faf1-UBX-domain binding and Ufd1-SHP motif binding to the same NTDs of p97 suggests a functional and physiological relevance.

### Faf1 promotes ubiquitin insertion and engagement by p97

To test whether the FH-UT3 domain interaction is critical for the Faf1-mediated stimulation of substrate unfolding through a potential acceleration of initiator-ubiquitin unfolding and insertion, we employed a FRET-based initiation assay that we previously developed for Cdc48-UN ^14^. For this assay, the second ubiquitin in unanchored long (> 10) ubiquitin chains was labeled on an engineered cysteine near its N-terminus with a fluorescent donor dye (Cy3). P97 subunits were labeled with an acceptor dye (LD655) on the unnatural amino acid azido-phenylalanine (AzF) that was introduced in place of D592, located near the exit of the central channel (Fig. 4A). Unfolding and threading of the donor-labeled ubiquitin is thus expected to bring it into proximity of the acceptor label and increase the apparent FRET efficiency. Control experiments confirmed that p97 with incorporated AzF and LD655 labels had unfolding activity, albeit at a reduced level compared to the unmodified enzyme (Supp. Fig. 11A-C). Measurements of fluorescence spectra after mixing the Cy3-labeled ubiquitin chains with ^LD655^p97-UN and ATP in the absence of Faf1 showed no difference in donor or acceptor fluorescence compared to the isolated components, whereas the presence Faf1 caused significant reciprocal changes in fluorescence intensities that are indicative of FRET (Fig. 4B). Faf1 thus appears to stimulate initiator-ubiquitin unfolding and insertion into the central channel, whereas UN alone is too inefficient for a considerable signal change over the time frame of this FRET assay, even though it can support slow substrate processing in our unfolding assay (Fig. 1C).

**Figure 4:**
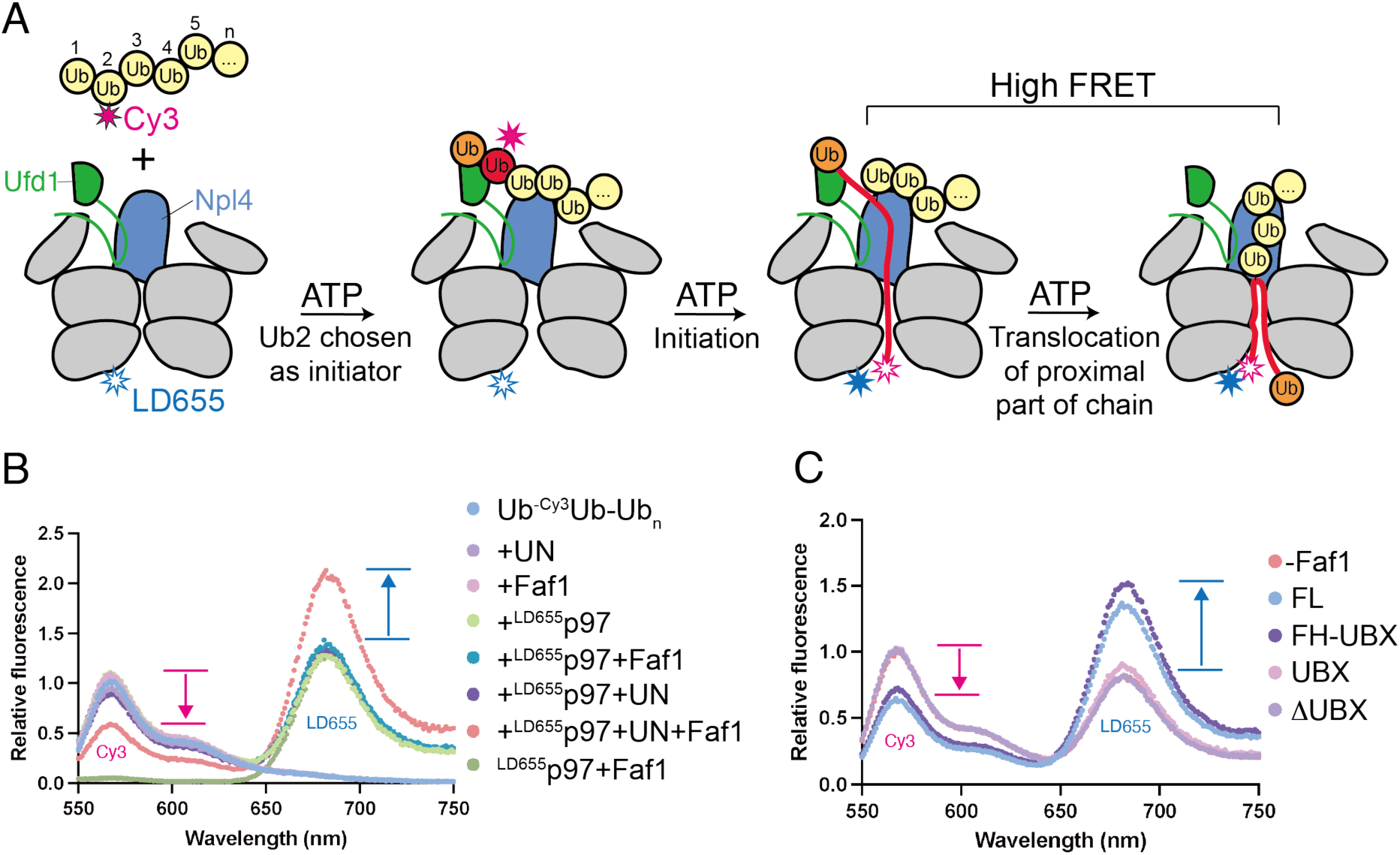
FRET-based unfolding-initiation assay. **A)** Experimental design for the initiation assay, using unanchored poly-ubiquitin chains (> 10 moieties) labeled with Cy3 (red star) on the N-terminus of the second ubiquitin and p97 labeled with LD655 (blue star) near the exit of the processing channel. UN choosing ubiquitin 2 as the initiator (top) leads to an immediate FRET-signal change after insertion of that ubiquitin moiety. **B)** Normalized fluorescence emission spectra after excitation at 480 nm for ubiquitin chains with Cy3 attached to ubiquitin 2 (Ub-^Cy3^Ub-Ub_n_) in the absence and presence of LD655-labeled p97 (^LD655^p97), UN, and Faf1, or for ^LD655^p97 with Faf1 alone. Arrows indicate the changes in Cy3– and LD655-fluorescence intensities upon unfolding initiation. **C)** Normalized fluorescence emission spectra after Cy3 excitation for samples with Ub-^Cy3^Ub-Ub_n_, ^LD655^p97, and UN in the absence or presence of different Faf1 truncation mutants. Arrows indicate the changes in Cy3– and LD655-fluorescence intensities upon unfolding initiation.

It is important to note that the changes in donor and acceptor signals were stable for an extended amount of time (> 10 min), suggesting that the ubiquitin chain either stalls in p97 or reaches a steady state of continuous unfolding and release, with the labeled second ubiquitin located inside or below the exit of the central channel and thus in proximity to the acceptor dye. For Cdc48-UN, initiator-ubiquitin unfolding and insertion was found to be fast (∼ 3 s) even in the presence of ATPγS ^14^, whereas p97-UN required ATP hydrolysis for a signal change in our initiation assay (Supp. Fig. 11D,E). It is possible that p97 depends on a combination of ATP– and ADP-bound subunits in the hexamer to adopt an appropriate spiral-staircase conformation for ubiquitin insertion. Alternatively, an initiator ubiquitin may be able to insert into ATPγS-bound p97, but the long chains necessary for efficient initiation by human p97-UN may strongly reduce the probability that the second, donor-labeled ubiquitin is chosen as the initiator in our assay. This model would also be consistent with our structural data, showing density for at least one additional ubiquitin besides Ub^prox^ bound to Ufd1’s UT3 domain. Initiation at position 3 or higher up in the chain would require ATP-hydrolysis-driven unfolding and translocation of proximal moieties to move the labeled ubiquitin 2 through the pore and into proximity of the acceptor for a FRET signal change (Supp. Fig. 11F).

Using the double mutations W241A/R242E or E446K/Y447A in the ubiquitin-binding groove of Npl4^12,14^, we confirmed that initiator-ubiquitin insertion into p97 still depended on its interaction with the Npl4 groove, even in the presence of Faf1 (Supp. Fig. 11G). In agreement with our findings for substrate unfolding, FRET-based initiation assays with truncated Faf1 variants showed that the UBX domain is critical for p97 binding and that the minimal FH-UBX domain fragment is sufficient for accelerating ubiquitin insertion into the central channel of p97 (Fig. 4C).

Despite the presence of Faf1, unfolding of the ubiquitinated Eos substrate by p97-UN is ∼ 200 times slower than by Cdc48-UN (Fig. 1C) ^14^. To assess potential differences in the kinetics of unfolding initiation, we monitored the FRET changes upon labeled ubiquitin insertion into p97-UN-Faf1 in stopped-flow mixing experiments. Interestingly, ubiquitin insertion occurred with an apparent time constant of 1_initiation_ = 5.1 ± 0.2 s (Supp. Fig. 11H), only ∼ 1.6-fold slower than for Cdc48-UN (1_initiation_ = 3.2 ± 0.1 s) ^14^. Notably, as described above, it is likely that a ubiquitin moiety distally located relative to the labeled ubiquitin 2 is chosen as the initiator, and additional unfolding and translocation of proximal ubiquitins is required for FRET signal development, such that the initiation time constant measured in our assay represent an upper bound. The velocity of substrate unfolding under single-turnover conditions is therefore not limited by slow initiation or the distinct ubiquitin-chain length requirements of p97-UN compared to Cdc48-UN, but due to slow Eos unfolding itself. To confirm that p97 does not get stuck on the folded Eos domain, we used an Eos-Turquoise fusion substrate. The observed loss in Turquoise fluorescence confirmed that Eos is completely unfolded and threaded by p97, albeit very slowly (Supp. Fig. 12). The significant difference in substrate-unfolding speeds for p97 and Cdc48 (1_Eos unfold_ = 901.5 ± 75.6 s and 1_Eos unfold_ = 4.4 ± 0.1 s, respectively) may originate from the almost two orders of magnitude different ATP-hydrolysis activities (16 min^-1^ for p97 versus ∼ 1000 min^-1^ for Cdc48 in the absence of UN and substrate ^14,20^). Furthermore, p97 may have extra layers of negative regulation that could be encoded through sequence differences or additional cofactors. Assuming that Faf1 primarily facilitates the unfolding and insertion of the initiator ubiquitin, we can use the observed time constants for Eos processing in the absence versus presence of Faf1 (1_Eos unfold_ (-Faf1) = 4727 ± 688 s and 1_Eos unfold_ (+Faf1) = 902 ± 76 s) together with the time constant for initiation with Faf1 from the FRET-based assay (1_initiation_ = 5.1 ± 0.2 s) to estimate that Faf1 accelerates initiation by more than 700 fold. Subsequent unfolding of the Eos moiety in our model substrate occurs much more slowly and becomes rate-limiting, accounting for ∼ 20% of the time spent on substrate processing in the absence of Faf1, which explains the maximum 5-fold overall acceleration by Faf1, despite its major effects on the efficiency and velocity of initiation.

### The stimulatory helix interaction with Ufd1’s UT3 domain is conserved for Faf2

Truncation of Faf1’s FH abolished Faf1’s stimulatory effects on substrate unfolding activity (Fig. 1F), consistent with our model that FH stabilizes Ufd1’s UT3 domain. Yet to further validate this hypothesis and our structural data for the ternary complex, we sought out point mutations in Faf1 and Ufd1 that sterically break the binding interface. Indeed, the S518K mutation in Faf1 or the A140E mutation in Ufd1 abolished the accelerated substrate unfolding by p97-UN-Faf1 (Fig. 5A,B; Supp. Fig. 15). Ufd1 A140E also reduced the photo-induced crosslinking between ubiquitin and the BPA incorporated at position 501 of Faf1 (Supp. Fig. 9), but had no effect on substrate unfolding in the absence of Faf1 (Fig. 5B).

**Figure 5:**
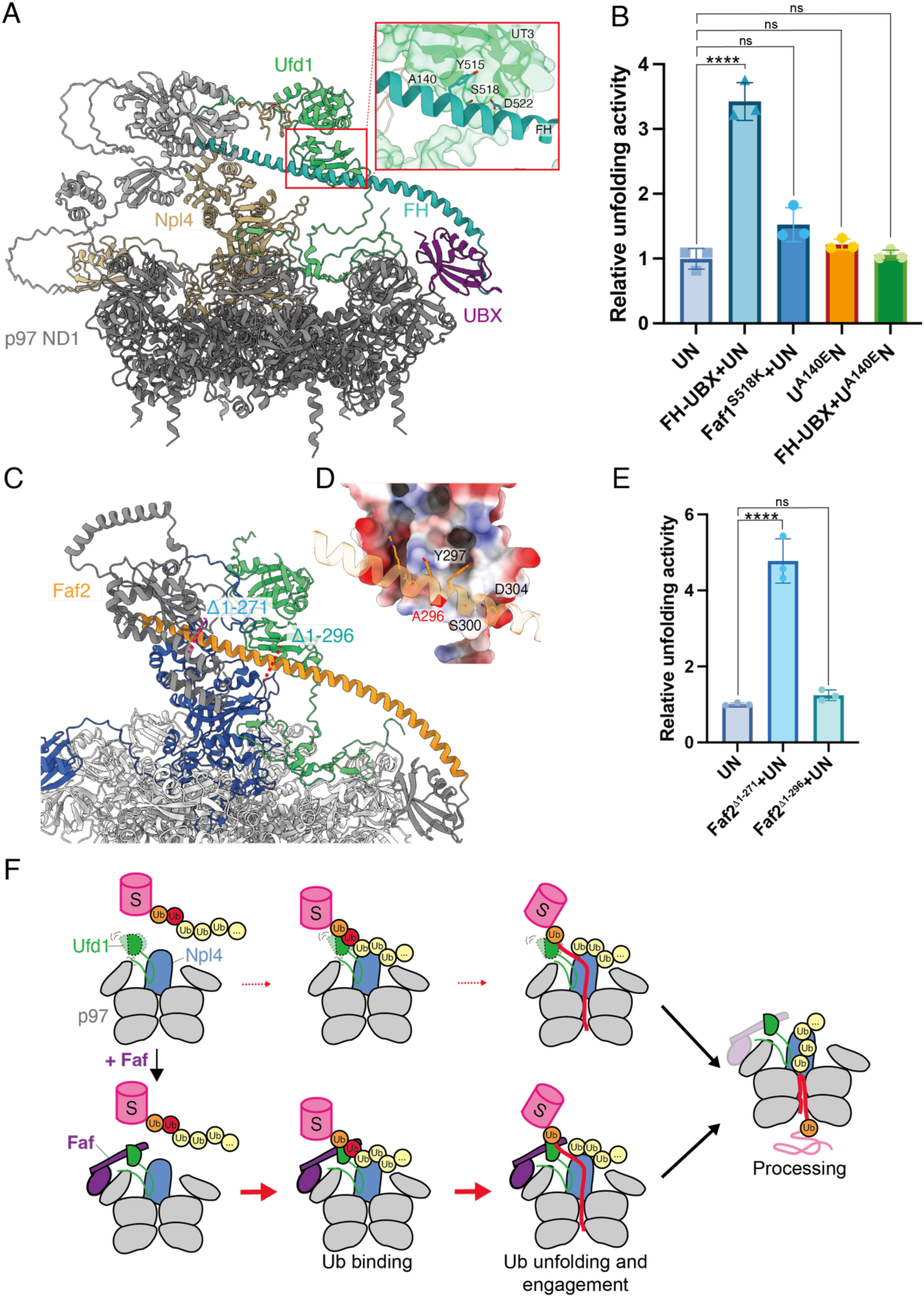
Faf1’s C-terminal helix facilitates initiator-ubiquitin unfolding and insertion into p97. **A)** AlphaFold3 model for the complex between p97’s N-terminal and D1 ATPase domains (ND1, dark grey), Npl4 (tan), Ufd1 (green), and Faf1, with Faf1’s UBA, UBL, and UAS domain depicted in light grey, the long helix (FH) in turquoise, and the UBX domain in purple. The red box shows a zoom-in of the interface between Faf1’s long helix and Ufd1’s UT3 domain. **B)** Relative mEos-substrate unfolding activities for p97 in the presence of wild-type UN, wild-type Faf1 FH-UBX fragment, UN with Ufd1 carrying an A140E mutation in its UT3 domain at the FH-binding interface, or Faf1 with a S518K mutation in its UT3-binding interface. Shown are the mean values and standard deviations of the mean for three technical replicates. Statistical significance was calculated using a one-way ANOVA test: ****p<0.0001; ns, p>0.05. **C)** AlphaFold 3 model for the complex between p97’s ND1 portion (light grey), Npl4 (dark blue), Ufd1 (green), and Faf2, with the long helix shown on orange and the rest of Faf2 in dark grey. Positions for Faf2 N-terminal truncation mutants Δ1-271 and Δ1-296 are indicated by red dashed lines. **D)** Close-up view of the interface between Ufd1’s UT3 domain and Faf2’s FH helix, with UT3 shown in surface representation and colored by electrostatic properties. **E)** Relative mEos-substrate unfolding activities for p97-UN in the absence or presence of Faf2 N-terminal truncation variants. Shown are the mean values and standard deviations of the mean for three technical replicates. Statistical significance was calculated using a one-way ANOVA test: ****p<0.0001; ns, p>0.05. **F)** Model for Faf-mediated engagement and unfolding of a ubiquitinated substrate (blue) by p97-UN. In the absence of a Faf cofactor, Ufd1’s UT3 domain may have higher flexibility that hampers the binding, unfolding, and motor engagement of the initiator ubiquitin (red sphere). Binding of Faf (purple) stabilizes the UT3 domain through its long helix that is anchored on its p97 NTD-bound UBX domain, facilitating unfolding initiation and overall substrate processing.

Faf2 is a membrane-associated cofactor that is involved in ERAD^28,29^, but shows partial conservation with Faf1 (Supp. Fig. 13). AlphaFold predicts that Faf2’s C-terminal UBX domain interacts with p97’s NTD, while the preceding FH binds to Ufd1’s UT3 domain (Fig. 5D). We generated two truncated Faf2 fragments, with a deletion of either the first 271 residues, which leaves the C-terminal helix and UBX domain, or 296 residues, which in addition removes about half of FH’s UT3-binding region and represents a fragment previously used in replisome-disassembly studies (Fig. 5D-E, Supp. Fig. 1D). In our Eos-unfolding assays, the addition of Faf2^Δ1-271^ to p97-UN led to a 5-fold stimulated activity (Fig. 5F), comparable to the FH-UBX domain fragment of Faf1. In contrast, Faf2^Δ1-296^ has some of the UT3-interacting residues removed and failed to accelerate substrate turnover. In summary, these results suggest that the helix preceding the UBX domain in both Faf1 and Faf2 interacts with the UT3 domain of Ufd1 to stimulate substrate processing by p97-UN, likely by stabilizing Ufd1’s position for more productive initiator-ubiquitin unfolding and insertion into the AAA+ motor.

## Discussion

Here we show that Faf1 and Faf2 accelerate the early steps in the processing of ubiquitinated substrates by p97-UN. The effects depend on the recruitment of these cofactors through their UBX domain to p97’s NTDs, which positions a conserved helix preceding the UBX domain to interact with the UT3 domain of Ufd1. This interaction appears critical for promoting a productive p97-UN complex that can rapidly insert the initiator ubiquitin into the ATPase motor, likely through stabilizing the position of Ufd1’s UT3 domain relative to Npl4. Our cryo-EM data together with AlphaFold modeling indicate that the UT3 domain binds not only the ubiquitin moiety (Ub^prox^) proximally neighboring the initiator ubiquitin (Ub^ini^), but also the C-terminus of Ub^ini^ itself, with the latter interaction apparently dependent on Faf1 bracing the UT3 domain. UT3-domain stabilization by Faf1 may thus facilitate the recruitment of the initiator ubiquitin, its unfolding, and its placement in Npl4’s hydrophobic groove for p97 insertion. In addition to a reasonably well-resolved Faf1 moiety that seems to stably interact with the UT3 domain, we observed density for a second Faf1, whose resolution for the long helix preceding the UBX domain depended on the position of the ubiquitin-bound UT3 domain. The two Faf1 copies bind to neighboring NTDs of p97 that are also occupied by the two SHP motifs of Ufd1, suggesting a cooperativity in these interactions and a possible double-sided stabilization of the UT3 domain by two Faf1 molecules.

In the presence of Faf1, we observed ∼ 5-fold faster kinetics of substrate processing by p97-UN. The extent of this acceleration is thereby limited by the slow unfolding of our Eos model substrate, while ubiquitin unfolding and p97 insertion themselves are accelerated by more than two orders of magnitude. Initiation by p97-UN in the absence of Faf1 is apparently highly inefficient. Unlike Cdc48-UN, which shows maximum initiation and substrate-unfolding rates with ubiquitin chains of 6 moieties and reasonable activities even with dimeric chains^14^, p97-UN requires much longer ubiquitin chains for any detectable unfolding activity^19^. Our findings that Faf1 strongly facilitates ubiquitin engagement thus explain previous reports about Faf1 lowering the chain-length threshold for p97-UN^19^. Directly bridging p97’s NTD and Ufd1’s UT3 domain through interactions with Faf1’s UBX domain and rigid helix thereby seem essential, and it is conceivable that other UBX-domain containing proteins use similar multivalent contacts to help coordinate the myriad of p97 cofactors. Interestingly, Faf1 contains several other domains, including a ubiquitin-binding UBA domain^26^ whose truncation did not cause significant defects in our unfolding experiments. Faf1 may thus engage in other substrate-specific interactions^23,27^ or even function in other pathways^32,40^.

Initiation by p97-UN in the presence of Faf1 occurs similarly fast as by Cdc48-UN, yet unfolding the Eos moiety of our model substrate takes p97 ∼ 200 fold longer than Cdc48. This is likely caused by the vastly different rates in ATP hydrolysis for these human and yeast orthologs. Previous studies of GFP degradation by the bacterial AAA+ protease ClpXP showed that successful unfolding of the beta-barrel structure critically depends on the pulling frequency of the motor^41^, such that p97 may have a considerable disadvantage in unraveling Eos and potentially have to compete with the fast refolding of a short-lived unfolding intermediate. Future studies will have to address whether p97 could benefit in its substrate-unfolding velocity from other cofactors that either stimulate ATP-hydrolysis or act as chaperones to destabilize the folded domains of substrate proteins.

## Resource availability

### Lead contact

Further information and requests for resources and reagents should be directed to and will be fulfilled by the lead contact, Andreas Martin.

## Materials availability

All constructs generated in this study are available from the lead contact upon request and completion of a Material Transfer Agreement.

## Data and Code Availability

- All data generated or analyzed during this study are included in this manuscript and the Supplemental materials. The cryo-EM density maps and corresponding atomic coordinates can be found on the Electron Microscopy Data Bank (EMDB) and the Protein Data Bank (PDB) with the following accession code: p97-UN-Faf1^FL^ EMD-73536, PDB-ID 9YW2; p97-UN-Faf1^FH-UBX^ (PI state) EMD-76074, PDB-ID 11VE; p97-UN-Faf1^FH-UBX^ (I state) EMD-76028, PDB-ID 11TA; p97-UN-Faf1^FH-UBX^ (Npl4 local) EMD-76026, PDB-ID 11SY; p97-UN-Faf1^FH-UBX^ (IC1 state) EMD-76053; p97-UN-Faf1^FH-UBX^ (IC2 state) EMD-76054.
- This paper does not report original code.

Any additional information required to reanalyze the data reported in this paper is available from the lead contact upon request.

## Acknowledgements

We thank the members of the Martin lab for helpful discussions. We also thank Rui Yan, Shixin Yang, and Adamo Mancino at the HHMI Janelia CryoEM Facility for help in microscope operation and data collection.

## Author Contributions

Conceptualization, Z.L. and A.M.; Methodology, Z.L., C.A., and A.M.; Investigation, Z.L. and C.A.; Writing – Original Draft, Z.L. and A.M.; Writing – Review & Editing, Z.L., A.M., and C.A.; Funding Acquisition, A.M.; Supervision, A.M.

## Methods

### Purification of recombinant proteins

Expression, purification, and labeling of p97 and p97^D592AzF^ were carried out as previously described^20,42^. Briefly, after the recombinant expression of wild-type p97 in *E. coli*, cells were harvested by centrifugation at 3,500 rcf and resuspended in NiA buffer (50 mM HEPES-KOH, pH 7.6, 500 mM KCl, 20 mM imidazole) supplemented with benzonase nuclease, lysozyme, and protease inhibitors. Cells were lysed by sonication, the lysate was clarified by centrifugation at 27,000 rcf, and the supernatant was loaded onto Ni-NTA resin followed by 10 column volumes (CV) washing with NiA buffer. The protein was eluted with NiB buffer (50 mM HEPES, pH 7.6, 500 mM KCl, 400 mM imidazole) and concentrated before loading on a size-exclusion chromatography column (HiLoad Superdex 200 16/60) that was equilibrated with GF buffer (50 mM HEPES-KOH, pH 7.6 150 mM KCl, 5 mM MgCl_2_, 1 mM DTT). Fractions were analyzed by SDS-PAGE, concentrated, and snap frozen by liquid nitrogen. For p97^D592AzF^ expression, cells were co-transformed with the plasmids coding for p97^D592AzF^ and the AzF-tRNA synthetase, grown at 37°C to an OD_600_ of 0.6, and incubated with AzF at a final concentration of 1 mM. After 30 min, expression was induced with 0.5 mM IPTG and cells were grown at 18°C for 16 h before harvest.

Expression and purification of human UN complex and mutans were carried our as previously described^20^. Briefly, *E. coli* BL21 DE3 cells with a plasmid coding for Ufd1 or Npl4 were grown separately and resuspended in NiA buffer together before cell lysis. The supernatant after centrifugation was subjected to Ni-NTA purification and UN was further purified by size-exclusion chromatography (HiLoad Superdex 200 16/60). Peak fractions containing the UN complex were pooled and concentrated before snap freezing.

Faf1, Faf2, and their variants were expressed in Rosetta2 pLysS cells as GST fusions with an HRV-3C protease cleavage sites. Cells were lysed, clarified by centrifugation, and loaded on a glutathione resin equilibrated with GF buffer. After washing with 20 CV of GF buffer, GST-HRV3C protease was added to the resin and incubated at 4°C overnight to cleave off the tag. Cleaved target proteins were washed off from the resin with GF buffer supplemented with 5 mM DTT and further purified by size exclusion chromatography (Superdex 200 increase 10/300).

For expression of Faf1 FH-UBX^R501BPA,E527C^, cells were transformed with a plasmid coding for Faf1 FH-UBX^E527C^ carrying an amber stop codon at residue 501 and the vector encoding the BPA aminoacyl-tRNA synthase (MjTyrRS) / tRNA pair for BPA incorporation^43,44^. Cells were grown at 37°C to an OD_600_ of 0.6, and incubated with pBPA dissolved in NaOH at a final concentration of 2 mM. After 30 min, expression was induced with 0.5 mM IPTG and cells were grown at 18°C for 16 h before harvest.

Wild-type human ubiquitin and its variants were purified as previously reported^14^. In brief, for wild-type ubiquitin and ubiquitin with a N-terminal Met-Cys extension (MC-Ub), cells were lysed in NiA buffer, the supernatant was clarified by centrifugation at 27,000 rcf for 30 min, and glacial acetic acid was gradually added until the solution reached a pH of ∼ 4.5. Precipitants were separated by centrifugation at 27,000 rcf for 30 min, the supernatant was dialyzed at 4°C for 16 h against 50 mM sodium acetate buffer, pH 4.5, and additional precipitants were removed by centrifugation. The supernatant was loaded to a cation-exchange column (5 mM HiTrap SP HP, Cytiva) equilibrated with 50 mM sodium acetate buffer, pH 4.5, and proteins were eluted with a linear gradient of 1-60 % of 1 M NaCl over 30 CV. Peak fractions containing the target protein were pooled and concentrated, and proteins were further purified by size exclusion chromatography (HiLoad Superdex 75 16/60) in GF buffer. For Ub-His and MC-Ub-His, a Ni-NTA purification step was performed instead of the acidic precipitation process. For all MC-Ub variants, 5 mM DTT was added to the GF buffer for the final size exclusion chromatography to keep cysteines reduced.

The plasmid for expression of His-SUMO-Ub^G76V^-mEos3.2 was synthesized by GenScript and used to transform BL21 Star (DE3) competent cells. Expression was induced at OD_600_ = 0.6, and cell were grown at 20°C for 16 h before harvest. After pelleting, cells were resuspended in NiA buffer and lysed by sonication. The lysate was cleared by centrifugation and the supernatant was loaded on Ni-NTA resin pre-equilibrated with NiA buffer. After washing with 20 CV NiA buffer, the target protein was eluted with NiB buffer. Imidazole in the solution was diluted using Amicon spin filters before the addition of SENP2 protease. The cleavage was performed at 4°C for 16 h, and cleaved Eos was isolated as the flow through fraction of a subtractive Ni-NTA purification. Samples were concentrated and further purified by size-exclusion chromatography (HiLoad Superdex 75 16/60).

For the fusion protein of Eos-Turquoise, BL21Star (DE3) competent cells were transformed with pET24 His-3C-Ub^G76V^-mEos3.2-Turqoise and grown as described above. The protein was purified by Ni-NTA affinity in NiA buffer and size-exclusion chromatography in GF buffer with the His tag uncleaved.

Mouse E1 (Ube1)^45^ and the E2-E3 chimera gp78RING-Ube2g2^20^ were expressed and purified as described.

### Protein labeling with fluorophores

After Ni-NTA affinity purification, p97^D592AzF^ was treated with 5-fold molar excess of DTNB for 30 min at room temperature to protect solvent-exposed cysteines and then labeled with 5-fold molar excess of LD655-DBCO (custom synthesis by Lumidyne Technologies) for 2 h at room temperature. The reaction was quenched with an excess amount of free AzF and treated with 5 mM DTT before a final purification by size exclusion chromatography (HiLoad Superdex 200 16/60) in GF buffer. The efficiency of labeling was estimated based on the protein concentration measured by BCA assay in comparison with LD655 absorbance (typically 20-30%).

MC-Ub and its variants were labeled with Cy3– or Cy5-maleimide dyes at the N-terminal engineered cysteine. First, MC-Ub was treated with 1 mM TCEP for 30 min at room temperature, followed by the addition of 10-fold molar excess of sulfo-Cy3-maleimide or sulfo-Cy5-maleimide (Lumiprobe) dissolved in DMSO. The final concentration of MC-Ub was 500 µM, and the labeling reactions were carried out at room temperature for 30 min. Reactions were quenched by adding DTT to a final concentration of 10 mM, the protein was concentrated and subjected to size-exclusion chromatography (Superdex 75 increase 10/300) in GF buffer. Protein concentration was estimated based on Cy3 or Cy5 absorbance. Faf1 FH-UBX^R501BPA,E527C^ was labeled in a similar way as MC-ubiquitins, since C527 is the only cysteine in this fragment.

### Preparation of ubiquitinated Eos and Eos-Turquoise substrates

The reaction mixture for ubiquitination consisted of 7.5 µM E1, 125 µM chimeric E2-E3, 100 µM Ub^G76V^-mEos3.2 or Ub^G76V^-mEos3.2-Turquoise, 500 µM ubiquitin and GF buffer with 10 mM ATP. The reaction was carried out in a total volume of 500 µL at 37°C for 2.5 h. An extra 500 µM ubiquitin was added every 30 min to efficiently elongate the ubiquitin chain. The mixture was subsequently subjected to size-exclusion chromatography (Superose 6 increase 10/300), and fractions with substrate-attached ubiquitin chain of more than 10 Ub were pooled and concentrated. To prepare the red Eos, purified ubiquitinated green Eos was photoconverted on ice for 30 min using a 395nm 50W LED Black Light lamp (CONALUX). The conversion efficiency was ∼ 50-60% based on the fluorescence ratio.

### Assembly of ubiquitin chains

To prepare Ub-^Cy3^Ub-Ub_n_ and ^Cy5^Ub-^Cy3^Ub-Ub_n_ chains used for FRET-based experiments, the first step was to make Ub-His-^Cy3^Ub and ^Cy5^Ub-His-^Cy3^Ub ^16^. Ub-His or ^Cy5^Ub-His was mixed with a limited amount of ^Cy3^Ub in the presence of the ubiquitination machinery in reaction buffer (50 mM HEPES-KOH, pH 7.6, 150 mM KCl, 5 mM MgCl_2_, 0.5 mM DTT and 10 mM ATP) at final concentrations of 1.5 μM Ube1, 25 μM gp78RING-Ube2g2, 250 μM Ub-His or ^Cy5^Ub-His, and 50 μM ^Cy3^Ub. Reactions were incubated at 37°C for 40 min before loading them onto Ni-NTA resin equilibrated with NiA buffer. Proteins were eluted with NiB buffer, concentrated, and subjected to size-exclusion chromatography (Superdex 75 increase 10/300) in GF buffer. Peak fractions were analyzed by SDS-PAGE, and fractions containing di-Ub were pooled and concentrated. Ub-His-^Cy3^Ub or ^Cy5^Ub-His-^Cy3^Ub were elongated with unlabeled ubiquitin in the presence of ubiquitination machinery at 37°C for 2.5 h. Final concentrations were 7.5 μM Ube1, 125 μM gp78RING-Ube2g2, 100 μM Ub-His-^Cy3^Ub or ^Cy5^Ub-His-^Cy3^Ub, and 1 M ubiquitin, with ubiquitin being added in 200 mM aliquots every 30 min. Ubiquitin chains were purified by Ni-NTA affinity and size-exclusion chromatography (Superdex 200 increase 10/300). Fractions with chains containing more than 8 Ub moieties were pooled and concentrated.

### Single-turnover unfolding assays

Single-turnover unfolding assays for ubiquitinated Eos or the Eos-Turquoise fusion substrate were performed using a BMG Labtech CLARIOstar plate reader at 37°C. Substrate was preincubated with ATP regeneration mix (see below), and p97 was incubated with cofactor(s) in assay buffer (50 mM HEPES-KOH, 150 mM KCl, 5 mM MgCl_2_, 1 mM DTT). The reaction was initiated by mixing the substrate with p97/cofactors in 1:1 volume ratio. The final concentrations were 20 nM substrate, 400 nM p97, 2 μM UN and/or 2 μM Faf, and 1X ATP regeneration mix (5 mM ATP, 0.03 mg/mL creatine kinase, 16 mM creatine phosphate). Unfolding was monitored by measuring the loss of fluorescence, at 585 nm after excitation at 565 nm for red Eos, at 520 nm after excitation at 500 nm for green Eos, and at 475 nm after excitation at 415 nm for Turquoise. Fluorescence traces were fitted by single-exponential decay functions using GraphPad Prism.

### Complex assembly and cryo-EM sample preparation

#### Sample for dataset 0

p97 was pre-incubated with UN, full-length Faf1, and Eos substrate carrying ubiquitin chains with ∼ 8 Ub moieties at 37°C for 10 min, before addition of 2 mM ATP and 10 more minutes of incubation. The final concentrations were 10 µM p97, 20 µM UN, and 20 µM Faf1. The mixture was subjected to size-exclusion chromatography (Superose 6 increase 10/300) in GF buffer supplemented with 2 mM ATPγS. Peak fractions were concentrated, concentrations were determined with BCA assays, and samples were snap frozen. For grid preparation, 3 µL of 4 mg/mL assembled complex was applied to an UltrAuFoil grid (r1.2/1.3, 300 mesh) that was glow-discharged using PELCO easiGlow system (25 mA, 25 s), and grids were plunge-frozen in liquid ethane using a Vitrobot Mark IV (Thermo Fisher Scientific) set to 4°C and 100% humidity with 4 s blot time.

#### Sample for dataset 1 and 2

^LD655^p97^D592AzF^ was pre-incubated with UN and Faf1 FH-UBX, and Ub-^Cy3^Ub-Ub_n_ was diluted in GF buffer supplemented with 5 mM ATP, 0.06% CHAPS, and 0.04% octyl-β-D-glucopyranoside on ice. p97/UN/Faf1 was then mixed with ubiquitin chains/ATP, and immediately loaded onto a grid (Ultrathin carbon on Quantifoil, R2/2, glow discharged in easiGlow, 25 mA, 90 s), followed by plunge freezing using Vitrobot (4°C, 100% humidity, 2 s wait time, 3 s blot time). The final concentrations were 12.5 µM ^LD655^p97^D592AzF^, 10 µM UN, 10 µM Faf1 FH-UBX, and 20 µM Ub-^Cy3^Ub-Ub_n_.

### Cryo-EM data collection, single-particle analysis, and model building

Grids were screened on a 200 kV Talos Arctica microscope (Cal Cryo Facility at UC Berkeley) and data collection was performed at the HHMI Janelia Research Campus, using SerialEM^46^ on a 300 kV Titan Krios G4 equipped with SelectrisX energy filter (6eV slit width) and a Falcon4i camera. For dataset 0, 24,016 movies were recorded at a nominal magnification of ×165,000 (0.743 Å/pix) with a total dose of 50 e^-^/Å^2^ nd a defocus range of –0.8 to –1.6 µm. For dataset 1 and 2, 10,496 and 6,748 movies were recorded at a nominal magnification of ×105,000 (1.182 Å/pix) with a total dose of 50 e^-^/Å^2^ and a defocus range of –0.8 to –1.6 µm. Detailed data processing steps are shown in Supp. Fig. 3 and 5.

For datasets 0, multi-frame movies were imported into cryoSPARC Live for motion correction and CTF estimation, and downstream steps were carried out using cryoSPARC v4.6 ^47,48^. Particles picked with Blob picker were subjected to 2D classification and ab-initio reconstruction to get rid of junk particles. A subset of the particle stack was used as seeds for Topaz training, and the trained model was used to extract particles out of the whole dataset^49,50^. After extraction, the particles were subjected to 2D classification, ab-initio reconstruction, and heterogeneous refinement. The main class with p97 NTD in the “up” conformation was further refined with C6 symmetry, and the density of five p97 subunits was subtracted after symmetry expansion to leave out a single subunit for focused 3D classification and local refinement of the p97-NTD/Faf1-UBX region.

For dataset 1 and 2, images were pre-processed as those in dataset 0. After initial hetero-refinement, particles in the major class were subjected to 3D classification with a mask covering the region above the D2 domain of p97. Two classes with clear Npl4 density were chosen for further particle sorting. For the one with tilted Npl4 density (PI state), further classification did not improve the heterogeneity on top of the Npl4 tower, likely due to Npl4’s flexibility in this state, and we therefore performed a global refinement. For the other state with more stable Npl4 density (I state), we first aligned particles on the p97 motor (motor focused map), and noticed heterogeneity on top of Npl4. This motivated us to align particles on Npl4, followed by 3D classification with a Npl4-focused mask. One class showed clear density for three distal ubiquitins and was further locally refined as Npl4 local map. Several other classes showed potential heterogeneity around the first distal ubiquitin, Ub^dist1^, but the numbers of particles were not sufficient for further processing. Another unique feature in the I state map is the Ufd1 UT3 density between two Faf1 helices. We performed 3D classification with a focused mask covering Faf1 FH-UBX and UT3, and chose two of these classes for further global refinement (IC1 and IC2).

For model building, an Alphafold3-predicted structural model was docked into the local map, manually adjusted using ISOLDE^51^ in ChimeraX^52^, COOT^53^, and finally refined using PHENIX real space refinement^54^.

### ATPase assays

The rate of ATP hydrolysis was measured using a coupled assay that monitors the decrease in NADH absorbance^55^ in a plate reader at 37°C. p97/cofactors and the substrate/ATPase mix (5 mM ATP, 1 mM NADH, 7.5 mM phosphoenolpyruvate, 12 U/mL pyruvate kinase, 18 U/mL lactate dehydrogenase) were pre-incubated separately and mixed in 1:1 volume ratio right before the measurement. The absorbance of NADH at 340 nm was monitored, and ATPase rates were determined by linear regression. The final concentrations were 50 nM p97, 250 nM UN, 250 nM Faf variant, and 250 nM ubiquitinated Eos substrate.

### FRET-based pore-insertion and initiator-unfolding assays

Steady-state FRET assays were performed using a BMG Labtech CLARIOstar plate reader at 37°C. p97 was pre-incubated with UN/Faf1 in assay buffer, and labeled polyubiquitin chains were incubated separately in assay buffer supplemented with ATP or ATPγS for 10 min. p97/UN/Faf1 was mixed with substrate in a 1:1 volume ratio and transferred to a 384-well plate. The plate was incubated in the plate reader at 37°C for 1 min before scanning the fluorescence emission from 550 nm to 750 nm after excitation at 480 nm. The final concentrations were 200 nM ^LD655^p97, 2 µM UN, 2 µM Faf variant, and 200 nM Ub-^Cy3^Ub-Ub_n_ for pore insertion assays. For the initiator-unfolding assay, final concentrations were 2 µM p97, 2 µM UN, 2 µM Faf variant, and 200 nM ^Cy5^Ub-^Cy3^Ub-Ub_n_. The fluorescence intensities for each spectrum were normalized relative to the Cy3 emission maximum at 568 nm.

The kinetics of initiator insertion into p97 were measured in an Auto SF120 stopped-flow fluorometer (Kintek). The syringes were loaded with 145 µL of ^LD655^p97/UN/Faf1 and substrates/ATP, respectively, and incubated at 37°C. After loading, 7 shots of 35 µL each were fired, and the last three were analyzed. The final concentrations were 1 µM ^LD655^p97/UN/Faf1 and 0.1 µM Ub-^Cy3^Ub-Ub_n_ chains. An excitation wavelength of 480 nm was used, and the fluorescence of donor and acceptor dyes were measured simultaneously in different channels.

### Gel-based crosslinking assay

Purified components were diluted and mixed in GF buffer supplemented with 2 mM ATP on ice, followed by a UV exposure with a 365 nm UV lamp for 45 min. The final concentrations were 2 µM p97, 4 µM UN or U^A140E^N, 2 µM Faf1 FH-UBX^R501BPA,E527C^ and/or 4 µM Ub-^Cy3^Ub-Ub_n_. Samples after crosslinking were analyzed by SDS-PAGE and imaged for Cy5 and Cy3 fluorescence, before Coomassie staining.

### AlphaFold predictions

All AlphaFold predictions were carried out on the AlphaFold 3 server^56^. For p97-UN in complex with Faf1 and Faf2, six copies of p97 NTD-D1 fragment (residues 1-451) were used instead of full length p97 due to the token limit. For Ufd1’s UT3 domain in complex with ubiquitins and Faf1 FH, residues 20-192 of Ufd1 and 473-565 of Faf1 were used.

## Supplemental Figures

**Supplemental Figure 1:**
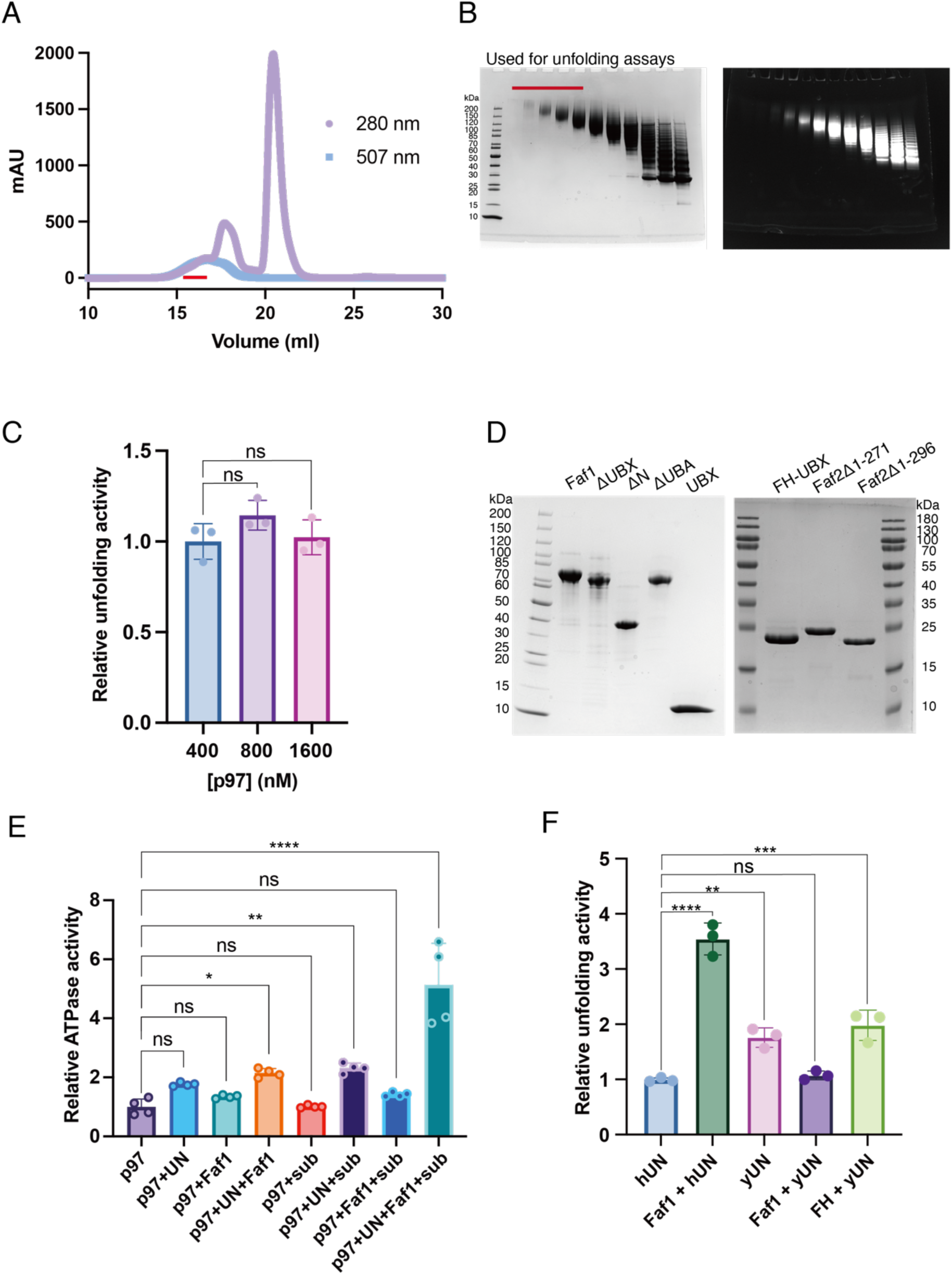
Sample preparations for biochemical experiments. **A)** Size-exclusion chromatogram for the purification of poly-ubiquitinated mEos substrate, monitored by general protein absorbance at 280 nm (purple) and mEos absorbance at 507 nm (light blue). Fractions pooled and used for unfolding experiments are indicated by a red bar. **B)** SDS-PAGE gels with the peak fractions of the size-exclusion chromatography shown in (A) and visualized by Coomassie staining (left) or mEos fluorescence (right). Fractions pooled and used for unfolding experiments are indicated by a red bar. **C)** Confirmation of single-turnover conditions for substrate unfolding. The rates for unfolding of the Eos substrate (20 nM) in the presence of UN (2 μM) and Faf1 (2 μM) did not significantly change when the p97 concentration was increased from 400 nM to 1600 nM, confirming saturating conditions for single-turnover kinetics. Shown are the mean values of the relative rates and the standard deviation of the mean for 3 technical replicates. Statistical significance was calculated using a one-way ANOVA test: ns, p>0.05. **D)** Coomassie-stained SDS-PAGE gels of purified full-length and truncated variants of Faf1 (left) and Faf2 (right). **E)** Relative ATPase activities of p97 in the absence (normalized to 1) and presence of UN, full-length Faf1, and ubiquitinated green Eos substrate. Shown are the mean values and standard deviations of the mean for four technical replicates. Statistical significance was calculated using a one-way ANOVA test: ****p<0.0001; **p<0.01; *p<0.05; ns, p>0.05. **F)** Relative rates for the unfolding of poly-ubiquitinated mEos substrate by p97 in the presence of human UN (hUN), yeast UN (yUN), and full-length Faf1 or the FH-UBX fragment. Shown are the mean values and standard deviations of the mean for three technical replicates. Statistical significance was calculated using a one-way ANOVA test: ****p<0.0001; ***p<0.001;**p<0.01; ns, p>0.05. The Eos substrate for these series of measurements carried slightly shorter ubiquitin chains (< 8 moieties) than the substrate for the measurements in Fig. 1F (8-12 ubiquitin moieties).

**Supplemental Figure 2:**
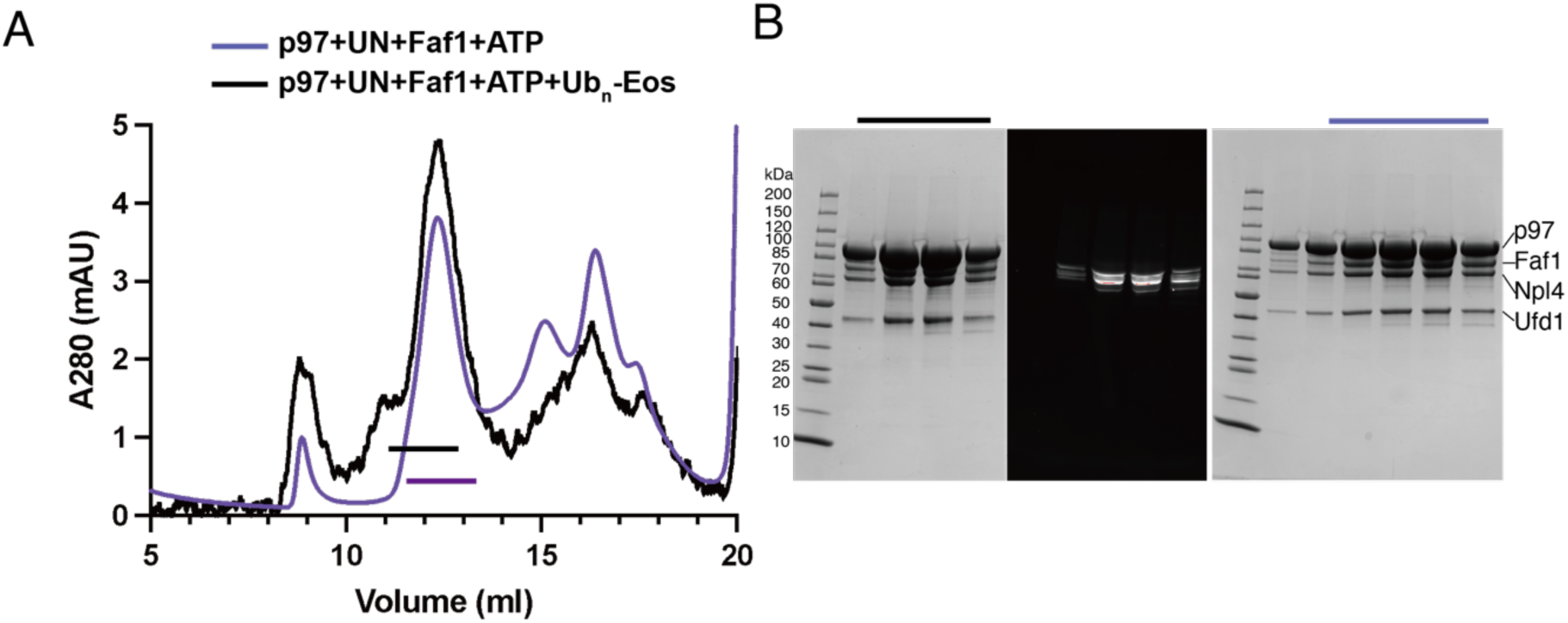
Sample preparation for cryo-EM structure determination of the p97-UN complex with full-length Faf1. **A)** Size-exclusion chromatogram for the purification of the p97-UN-Faf1 complex with ATP in the absence (purple) or presence (black) of poly-ubiquitinated (5-7 moieties) mEos substrate. Horizontal bars indicate fractions that were pooled and used for cryo-EM. **B)** SDS-PAGE gels with the peak fractions of the size-exclusion purifications shown in (A), visualized by Coomassie-staining (left) and mEos fluorescence (middle) for the substrate-containing sample, and visualized by Coomassie-staining for the substrate-free sample (right). Black and purple bars indicate the fractions that contained all components, were pooled, and used for cryo-EM sample preparations.

**Supplemental Figure 3:**
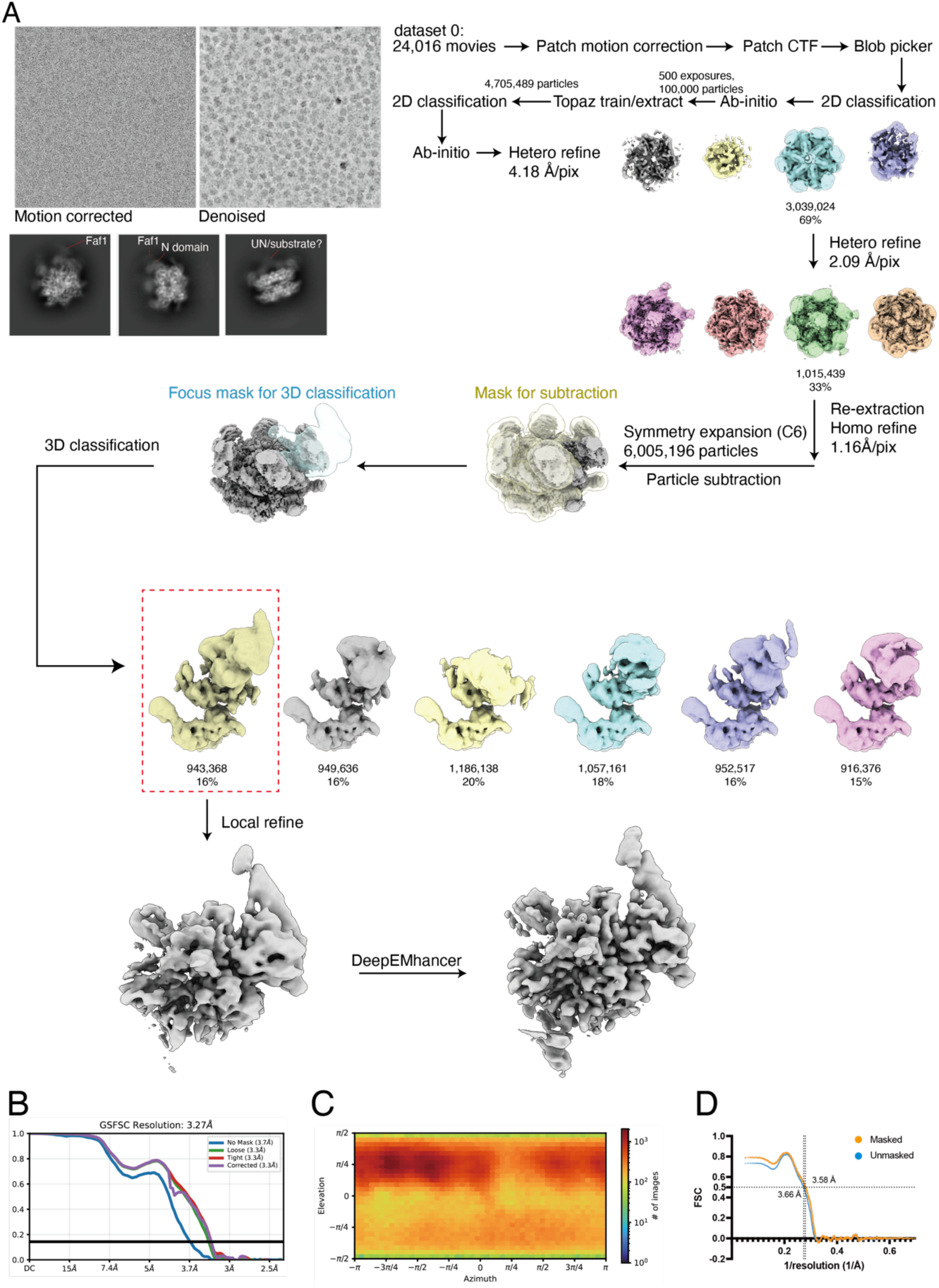
Cryo-EM single-particle analysis of p97-UN-Faf1 in complex with a ubiquitinated mEos-model substrate (dataset 0). **A)** Data-processing workflow using CryoSPARC. **B)** GSFSC curve and **C)** orientation distribution of the final map. **D)** Map-model FSC curve calculated after the final refinement in PHENIX.

**Supplemental Figure 4:**
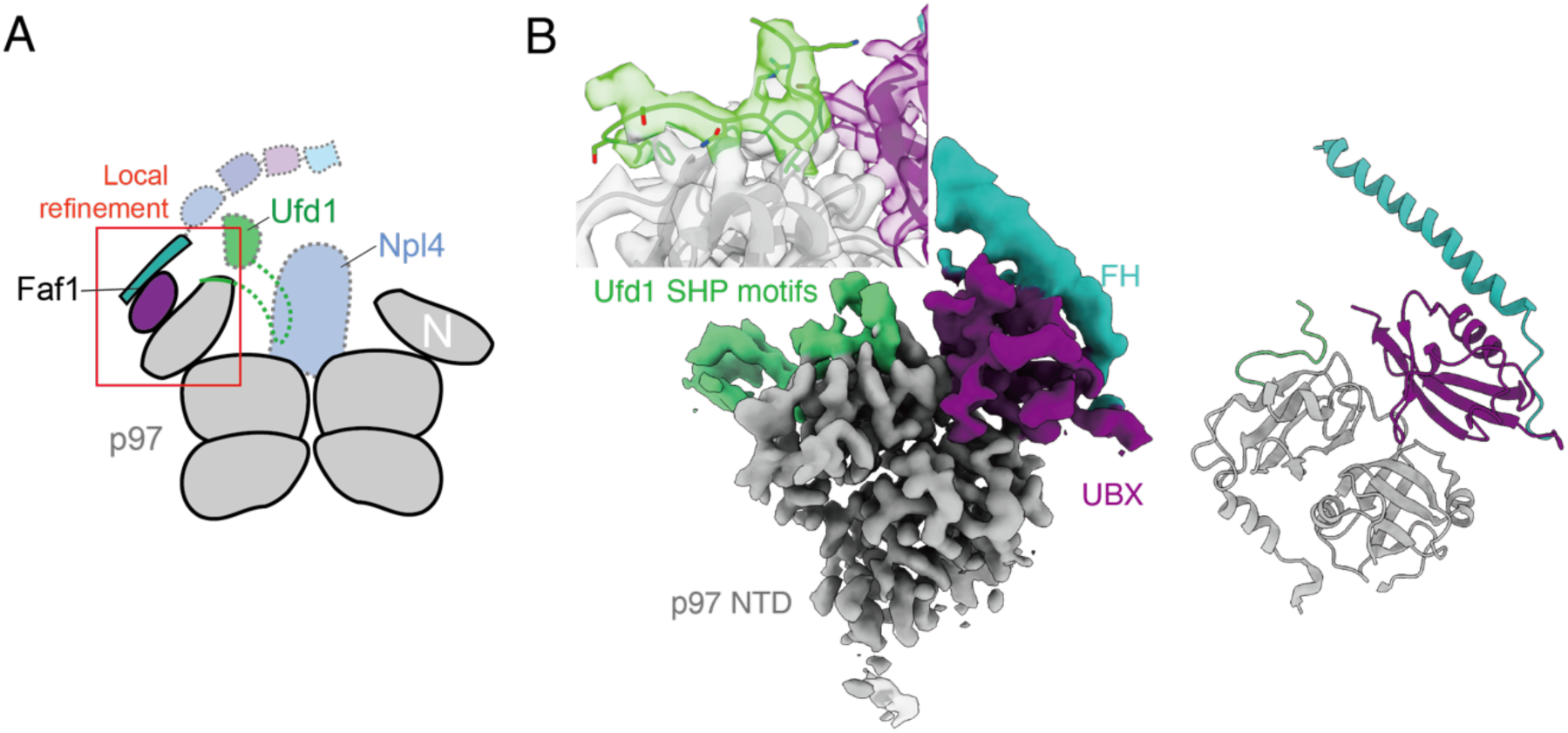
Cryo-EM structure determination of the p97-UN complex with full-length Faf1. **A)** Schematic of the p97-UN-Faf1 complex, with parts not resolved in this cryo-EM structure shown faded and with dashed outlines. The red box indicates the area for focused classification and local refinement in our single-particle analysis. **B)** Cryo-EM density (left) and atomic model (right) of the complex between p97 NTD (grey), Faf1’s helix (FH, turquoise) and UBX domain (purple), and Ufd1’s SHP motif (green). A focused view of the transparent map and atomic model of the SHP-interacting region is shown on the top left.

**Supplemental Figure 5:**
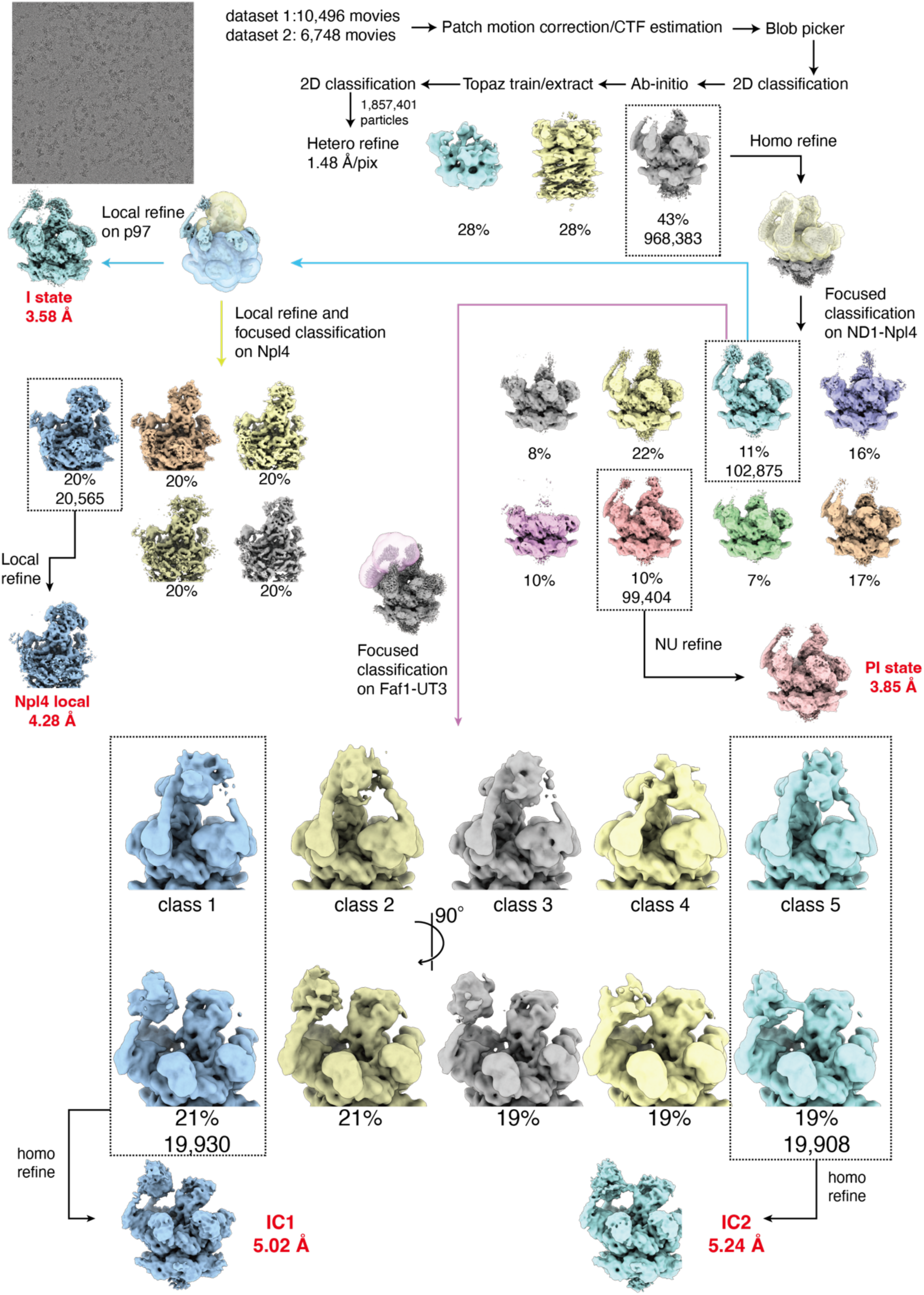
Cryo-EM single-particle analysis of p97-UN-Faf1^FH-UBX^ in complex with unanchored ubiquitin chains complex (dataset 1 and 2).

**Supplemental Figure 6:**
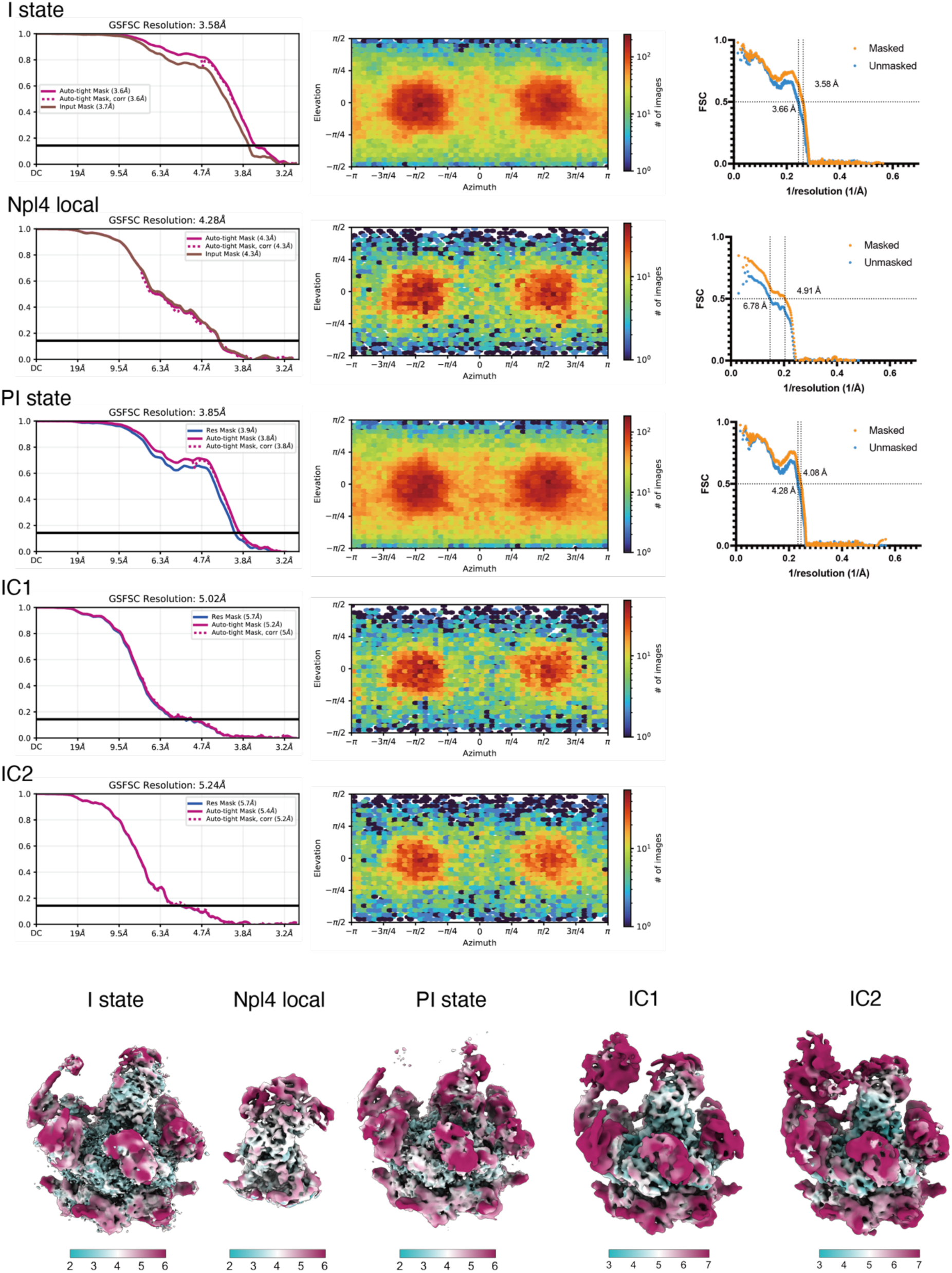
FSC curves, particle orientation distributions and local resolution maps for ubiquitin-chain-bound p97-UN-Faf1^FH-UBX^ ubiquitin bound complexes. The gold-standard Fourier shell correlation (GSFSC) curves are shown for each conformational state, reporting overall resolution at FSC = 0.143. Map-to-model curves are plotted for the PI state, I state, and the Npl4 local map. Local resolution maps are colored from high resolution (magenta/pink) to low resolution (cyan/teal) as indicated by the color bar beneath each map (Å).

**Supplemental Figure 7:**
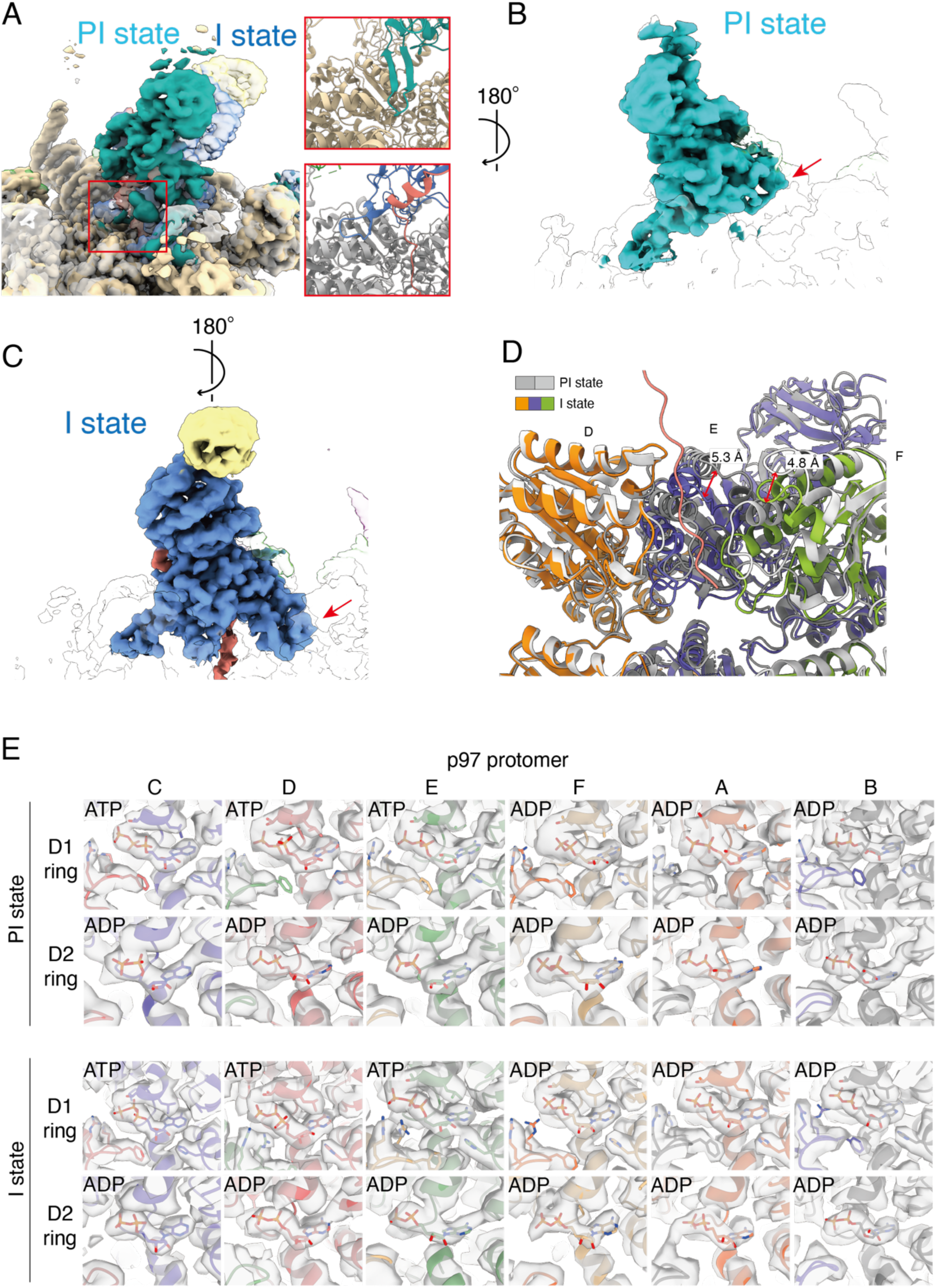
Comparisons of pre-initiation (PI) and initiation (I) states of the p97-UN-Faf^FH-UBX^ in complex with ubiquitin chains. **A)** Left: Overlay of the cryo-EM densities for the PI state (teal) and I state (blue), illustrating the conformational difference for Npl4. Right: Insets show zoomed views for the Npl4 loop (residues 425-439) that is positioned above p97’s processing channel in the PI state (top box), but relocated to allow insertion of Ub^ini^ (salmon) into p97 in the I state (bottom box). **B,C)** Density map of Npl4 in the PI state (B) and I state (C) is shown from a different angle to highlight the varied conformations of ZF2, indicated with red arrows. **D)** Overlay of the p97 D1 ATPase ring for the PI state (gray) and I state (colored) shows a conformational change of protomers E and F, which in the context of substrate engagement move downward toward the D2 ring. **E)** Cryo-EM densities and atomic models for individual nucleotide-binding pockets of all D1 and D2 ATPase domains in the PI and I states. Each panel is labelled with the identity of the bound nucleotide (ATP or ADP) that is shown in stick representation.

**Supplemental Figure 8:**
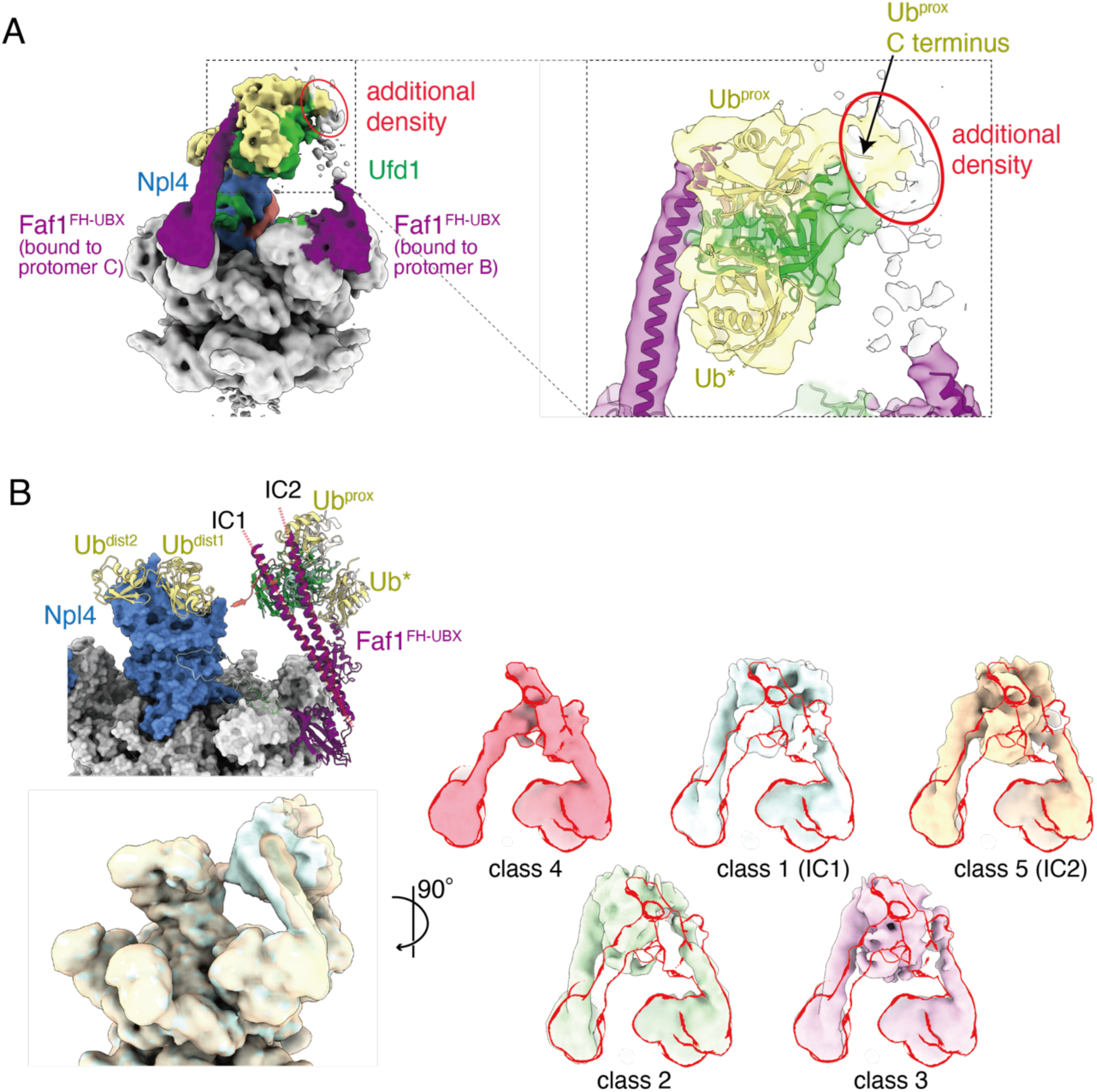
Ufd1’s UT3 domain is braced by a second Faf1 in a conformation-dependent manner. **A)** Side view of the cryo-EM density map for the IC1 state (left) and a closed-up view of the UT3-domain region with the docked-in atomic model for the ubiquitin-bound UT3 domain and Faf1 helices (right). **B)** Top left: Models for the IC1 and IC2 states are shown superimposed on the p97-Npl4 region (surface representation), revealing a movement of the ubiquitin-bound UT3 domain (dark colors for IC1, light colors for IC2) and the attached helix of the protomer-C-bound Faf1 away from Npl4 in IC2. Bottom left: Superimposed cryo-EM maps for the 3D classes of IC1 (cyan) and IC2 (yellow) particles. Right: Side views of the Faf1 FH-UT3 regions for five different classes overlayed with class 4 and in the same orientation as in panel A. These overlays highlight the cooperative motion of the UT3 domain and the two FH helices, with variable resolution for the second, protomer-B-bound Faf1.

**Supplemental Figure 9:**
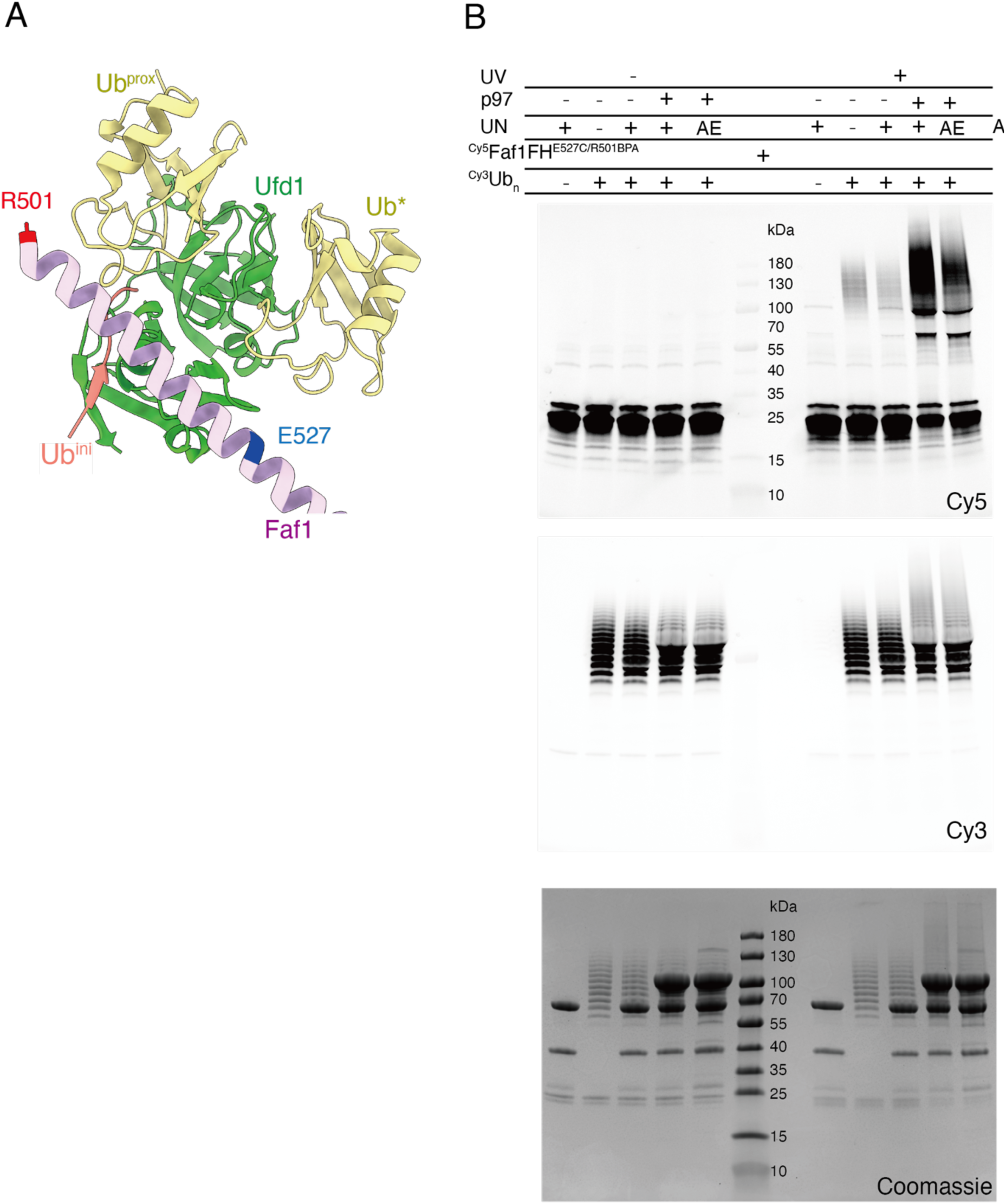
Gel-based assay for the photo-induced crosslinking between Faf1 FH and ubiquitin. **A)** IC1-based atomic model for the UT3-FH-ubiquitin complex to indicate the positions of the E527C mutation for Cy5 labeling and R501BPA incorporation for UV-induced crosslinking of Faf1 FH. **B)** SDS-PAGE gel images scanned for Cy5 (top) and Cy3 fluorescence (middle), and Coomassie-stained (bottom).

**Supplemental Figure 10:**
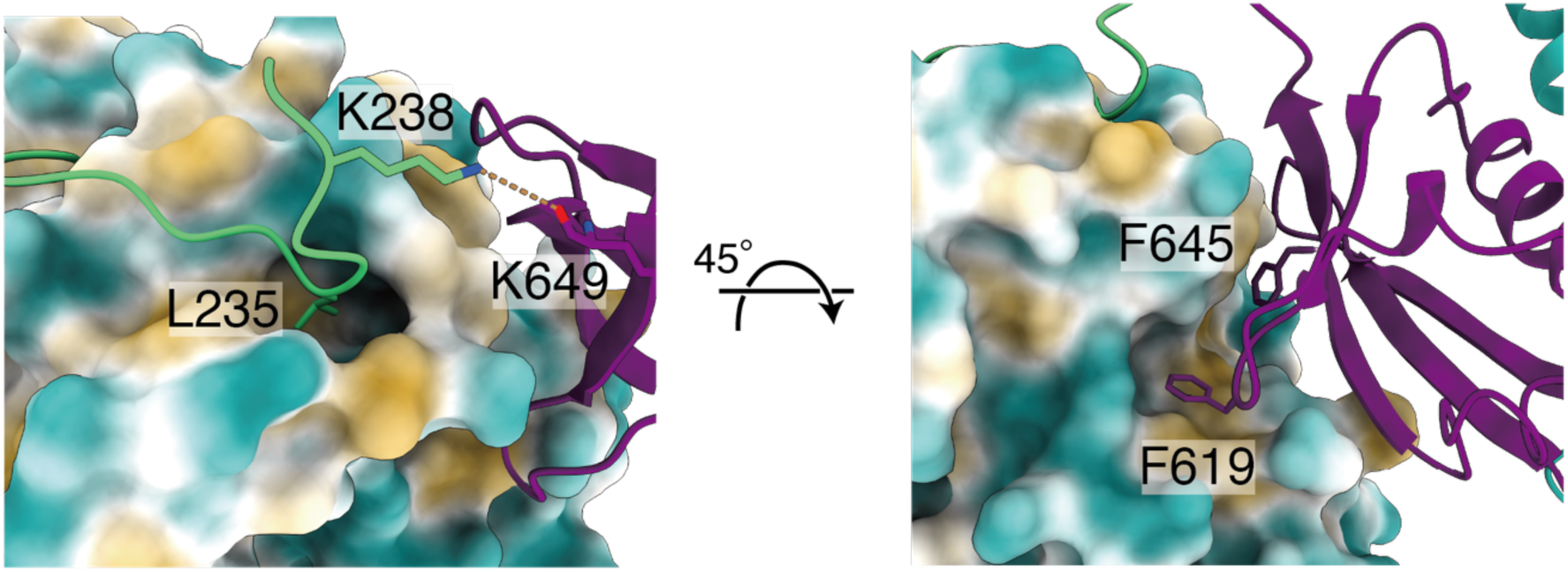
Close-up view of interactions between p97 NTD (represented as surface and colored by hydrophobicity), Ufd1’s SHP motif (green), and Faf1’s UBX domain (purple).

**Supplemental Figure 11:**
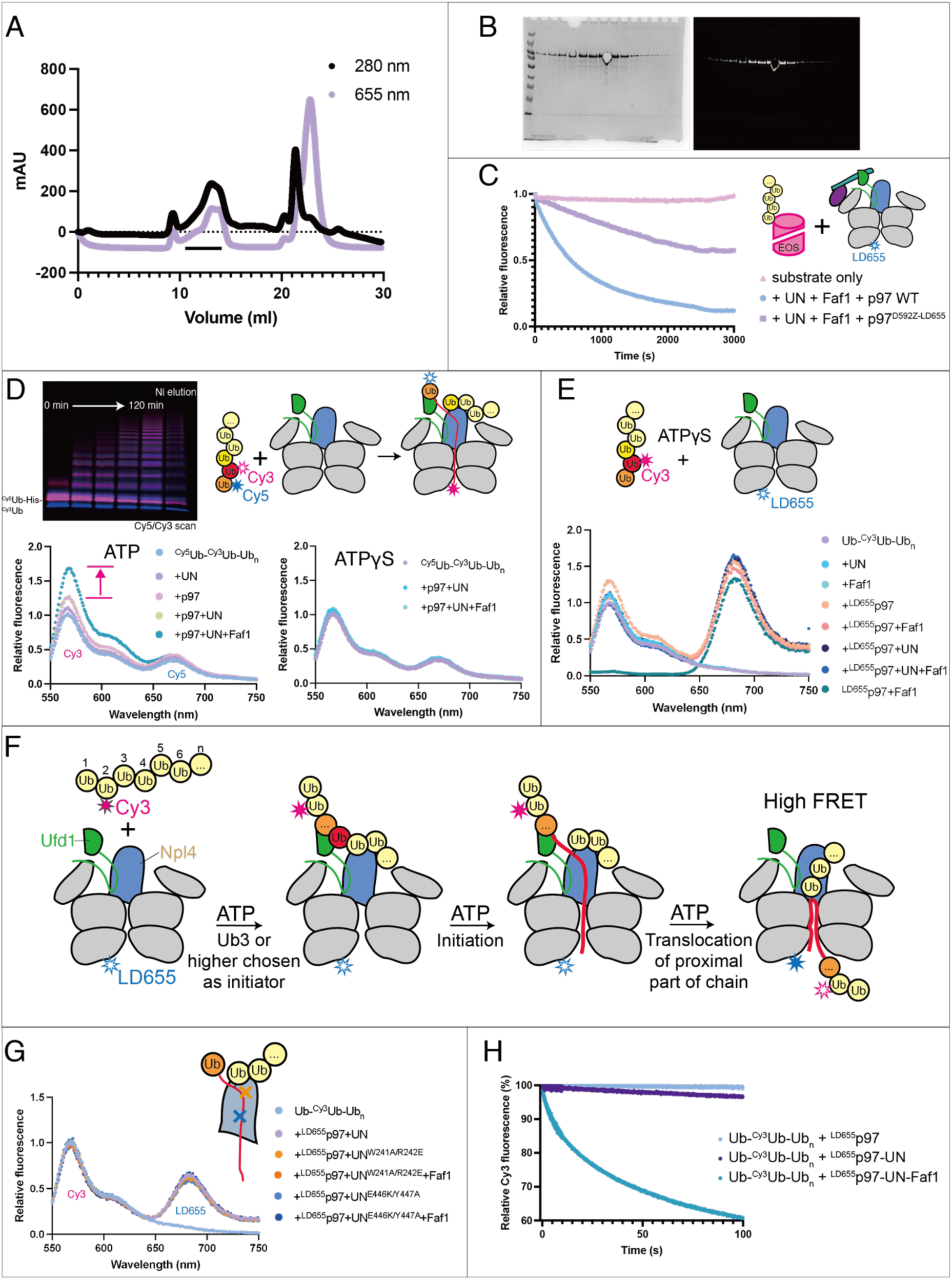
Extended results for FRET-based ubiquitin unfolding and initiation assays. **A)** Size –exclusion chromatogram for the purification of LD655-labeled p97^D592AzF^, detected by protein absorbance at 280 nm (black) or LD655 absorbance at 655 nm (purple). **B)** SDS-PAGE gels with the peak fractions of the purification show in (A) and visualized by Coomassie stain (left) or LD655-fluorescence scan (right). **C)** Example traces for the mEos-substrate unfolding by wild-type p97 or LD655-labeled p97, both in the presence of UN and Faf1 cofactors. **D)** Initiator-ubiquitin unfolding measured by the loss of FRET between Cy5 attached to the first ubiquitin and Cy3 attached to the second ubiquitin in unanchored ubiquitin chains (^Cy5^Ub-^Cy3^Ub-Ub_n_) after incubation with p97, UN, and Faf1. The gel image on the top left shows an overlay of Cy3– and Cy5-fluorescence scans for the SDS-PAGE analysis of doubly labeled ^Cy5^Ub-^Cy3^Ub-Ub_n_ synthesis progression over 120 min and the final sample after Ni-NTA affinity purification. Shown at the bottom are the normalized fluorescence emission spectra after excitation at 480 nm for the samples incubated with ATP (left) or ATPγS (right). **E)** FRET-based unfolding initiation assay with unanchored Ub-^Cy3^Ub-Ub_n_ ubiquitin chains, ^LD655^p97, UN, and Faf1 as shown in Fig. 4, but using ATPγS instead of ATP. Shown are normalized example spectra for the fluorescence emission after excitation at 480 nm. **F)** FRET-based initiation assay with unanchored Ub-^Cy3^Ub-Ub_n_ ubiquitin chains and ^LD655^p97 as shown in Fig.4A, but assuming that UN chooses ubiquitin 3 or higher in the chain as the initiator moiety, such that ATP-dependent unfolding and translocation of the proximally located ubiquitins is required to move the Cy3-labeled second ubiquitin into proximity of LD655 for a FRET-signal change. **G)** Same FRET-based initiation assay, but using Npl4 variants carrying W241A/R242E (orange) or E446K/Y447A double mutations (blue) in the hydrophobic groove that prevent capturing the unfolded initiator ubiquitin. Shown are the normalized fluorescence emission spectra after Cy3 excitation at 480 nm. **H)** Effects of Faf1 on the kinetics of unfolding initiation by p97-UN. Shown are example traces for the Cy3-fluorescence quenching after stopped-flow mixing of unanchored Ub-^Cy3^Ub-Ub_n_ ubiquitin chains with ^LD655^p97-UN in the absence or presence of Faf1.

**Supplemental Figure 12:**
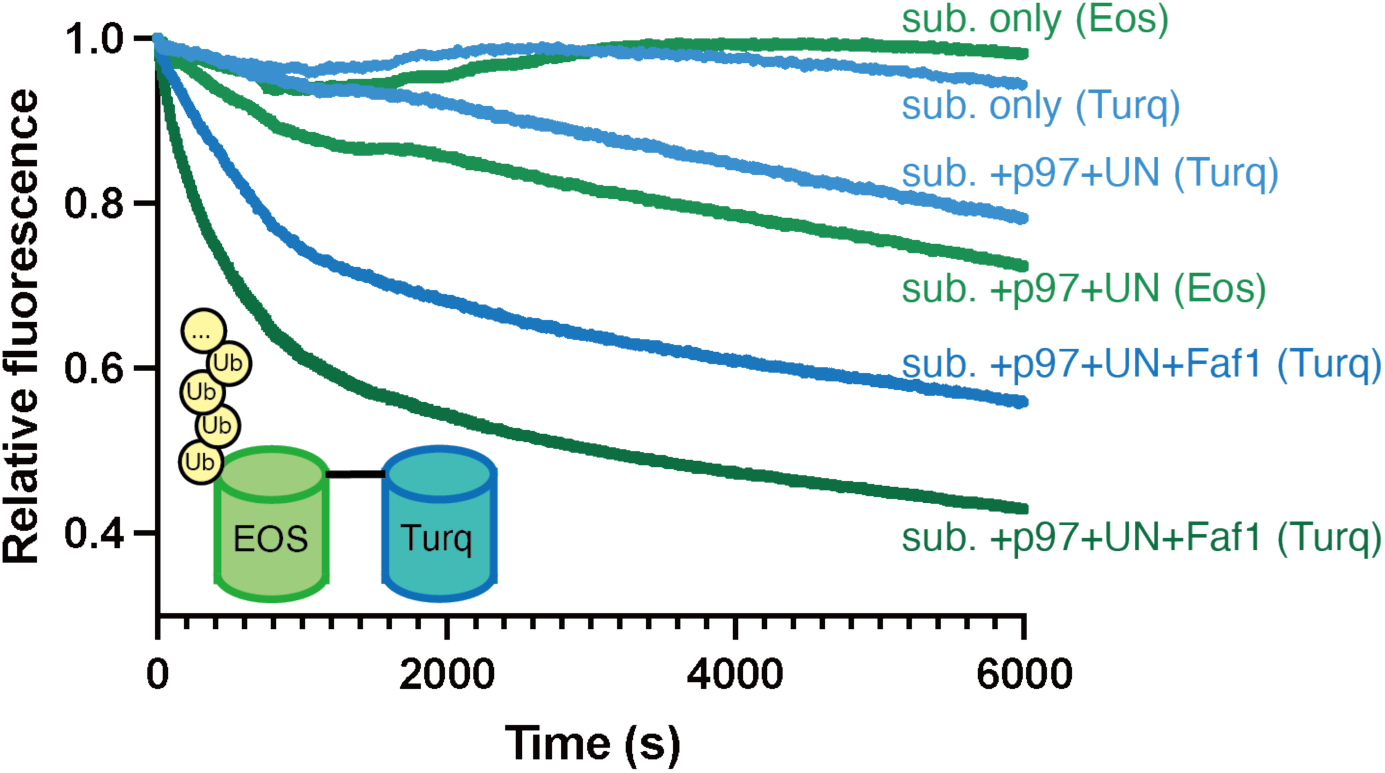
Kinetics for the unfolding of the poly-ubiquitinated mEos-Turquoise fusion substrate (schematic inserted in the bottom left) by p97-UN in the absence and presence of Faf1. Shown are example traces for the changes in green Eos fluorescence (shades of green) and changes in Turquoise fluorescence (shades of blue) that were measured separately after mixing the fusion substrate with p97 and cofactors. Loss in Turquoise fluorescence indicates that the Eos moiety of the substrate was completely unfolded and translocated.

**Supplemental Figure 13:**
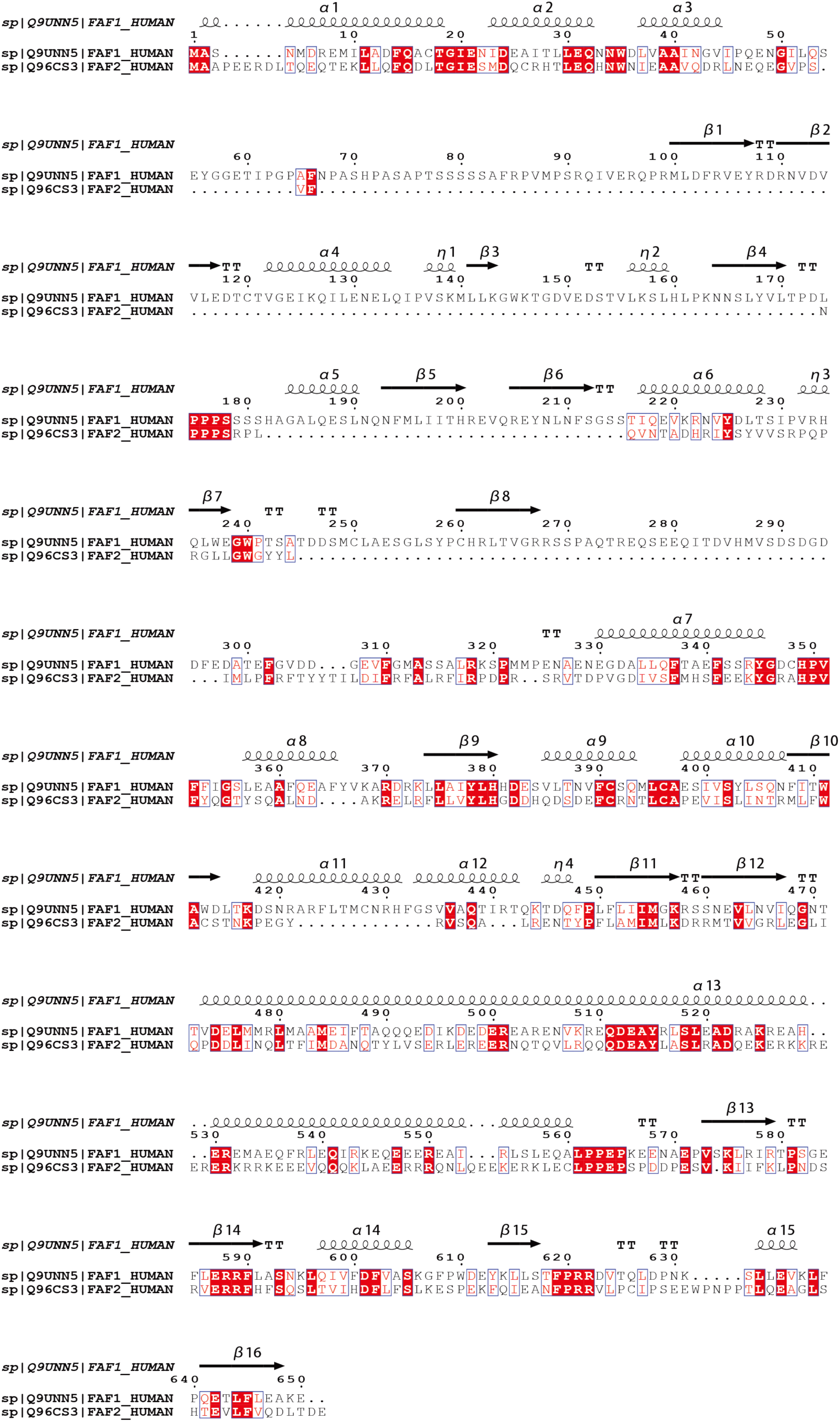
Sequence alignment of human Faf1 and Faf2.

**Supplemental Figure 14:**
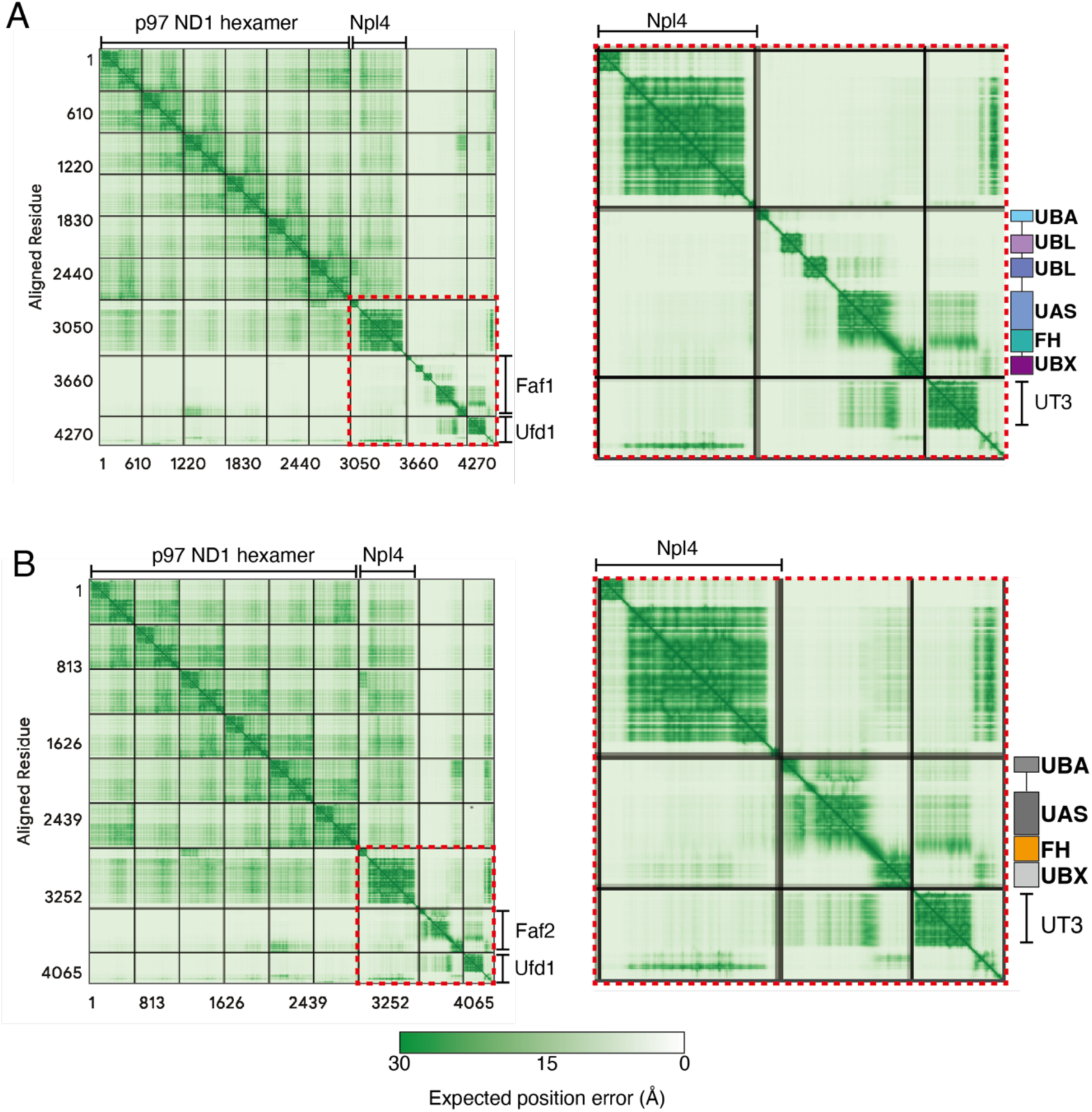
Predicted align error (PAE) for the AlphaFold 3 models of the p97-UN-Faf complexes. **A)** Left: PAE plot for the entire input of p97 NTD-D1, UN, and Faf1. Right: Zoom-in view of the Faf1-UN interaction, as highlighted by the red dashed box on the left. **B)** Left: PAE plot for the entire input of p97 NTD-D1, UN, and the Faf2 UBA-UAS-FH-UBX fragment. Right: Zoom-in view of the Faf2-UN interaction, as highlighted by the red dashed box on the left.

**Supplemental Figure 15:**
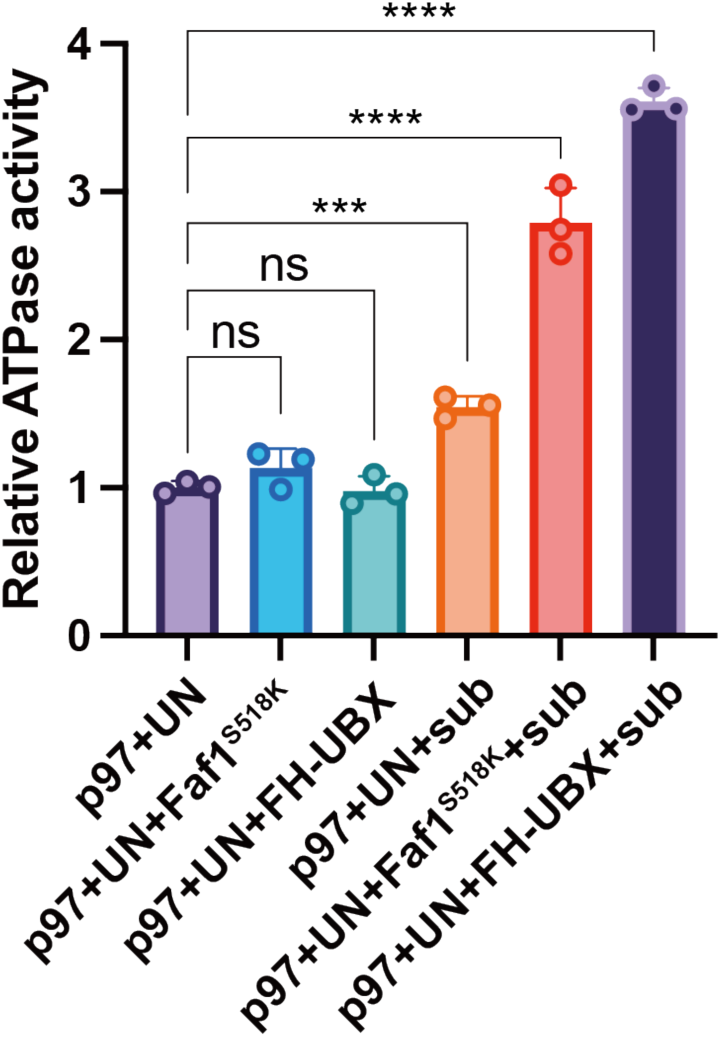
Relative ATPase activities of p97-UN (normalized to 1) and its complexes with Faf1^S518K^ of the Faf1 FH-UBX fragment, in the absence or presence of ubiquitinated green Eos substrate. Shown are the mean values and standard deviations of the mean for three technical replicates. Statistical significance was calculated using a one-way ANOVA test: ****p<0.0001; ***p<0.001; ns, p>0.05.

**Table 1:**
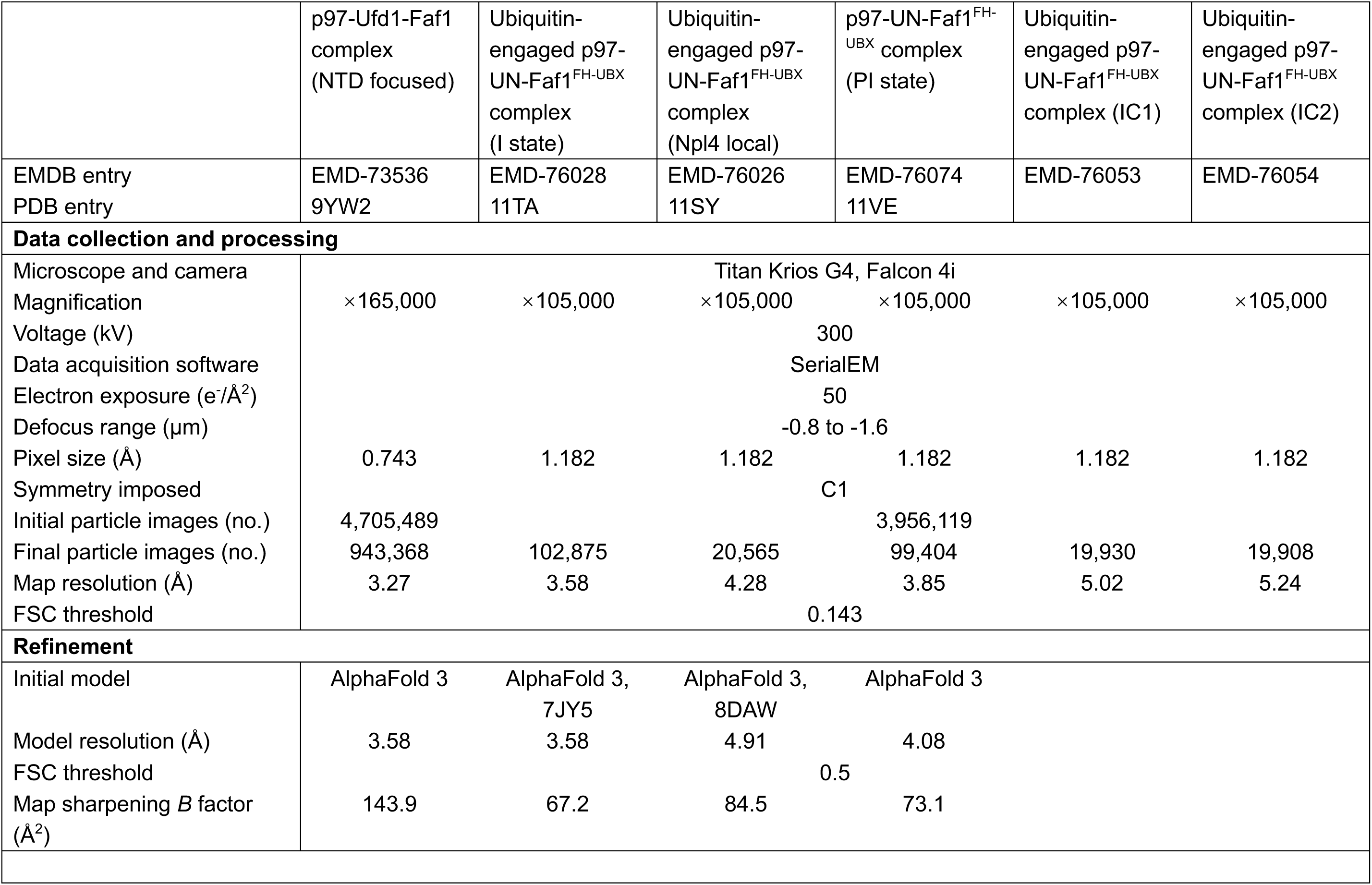
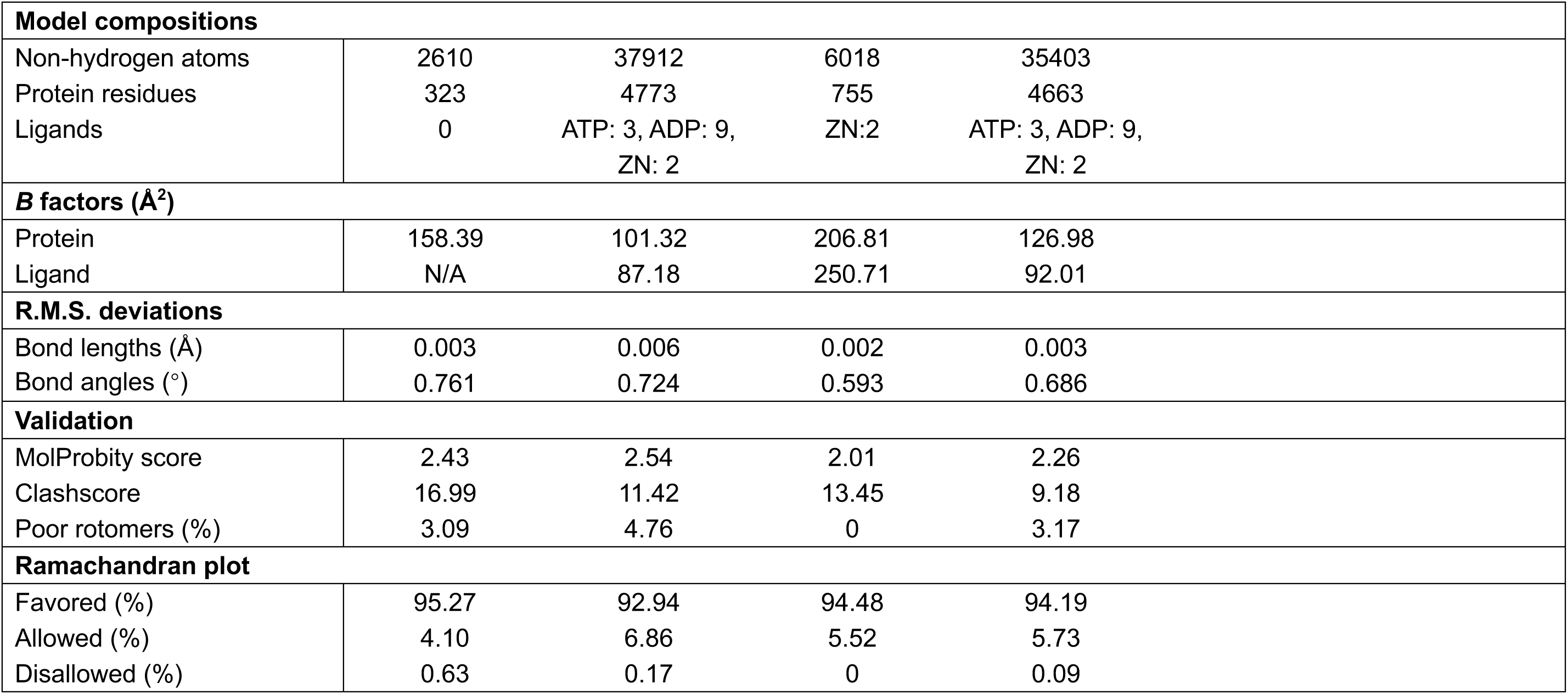
Cryo-EM data collections, refinement and validation statistics, and atomic models for the p97-Ufd1-Npl4 complex with full-length Faf1 and a ubiquitinated substrate or the Faf1 FH-UBX fragment in the presence of free ubiquitin chains.

